# Learning distributed representations with complementary components - A generative approximation framework for episodic memory encoding

**DOI:** 10.64898/2026.10.18.759853

**Authors:** Hao Liu, Marcelo Mattar, Sebastian Michelmann

## Abstract

Episodic memory must preserve detail while storing memories efficiently. We propose a framework of memory encoding in which generative neural network models approximate input through their frozen weights, yielding starting points from which the same weights can reinstate the input. This removes the need for mutually invertible cortical-hippocampal mappings, realizes a tradeoff between computational effort and mnemonic precision, and uses statistical regularities of the stimulus space to find representations that are distributed across the network. We implement this principle with a toy model and with large image-generation AI models, operationalizing mnemonic precision as closer approximation. Our model reproduces key findings from the Mnemonic Similarity Task: shallow encoding selectively impairs lure rejection, as observed in aging populations. Encoding further benefits from expertise: domain-finetuned models need fewer approximation steps for images from their own domain than for images from other domains. Finally, the representational geometry of latent starting points resembles medial temporal lobe but reinstatement through the weights of the network traverses the visual hierarchy in reverse order. Our framework offers a parsimonious and flexible account of memory encoding, where a generative search for a reinstatement-capable starting point finds a path for retrieval directly.

## Main

Episodic memory must balance two competing demands: preserving rich perceptual detail while still enabling efficient storage [1]. Both theoretical and empirical work suggest that memory precision improves with deeper processing and longer encoding times [2–4]. Moreover, human functional magnetic resonance imaging (fMRI) links hippocampal activity during memory encoding to the precision of subsequent retrieval [5]. The requirement to balance efficiency against detail consequently raises two fundamental questions: how the brain represents episodic memories in the first place, and how representations at different levels of detail are computed within its neural architecture.

Any answer to these questions is constrained by the architecture of human memory, which comprises two systems with different learning properties. The hippocampus is critical for episodic memory formation [6]; its neural activity lies on low-dimensional manifolds that jointly encode physical and abstract variables, forming geometric cognitive maps [7], and its neurons capture conceptual information in sparse codes [8–10]. Neocortical networks, on the other hand, learn slowly and gradually extract statistical regularities across many experiences [11–13].

Existing theories explain episodic retrieval as the reconstruction of distributed cortical activity from a hippocampal trace. The hippocampal indexing theory, for instance, proposes that the hippocampus stores pointers to distributed cortical patterns, enabling their reconstruction during recall [14]. Empirically, retrieval recruits a widespread recollection network comprising the hippocampus and content-independent cortical regions, together with content-specific cortical regions that reinstate patterns expressed during encoding [15–21]. Rolls [22, 23] proposes that polysynaptic back-projections from CA1 address cortical neurons to reinstate patterns, consistent with anatomical pathways through the subiculum and entorhinal cortex [24, 25].

Recent computational models have begun to formalize the role of cortical weights in supporting episodic memory. Spens and Burgess propose that one-shot hippocampal learning is complemented by a generative cortical network that reconstructs sensory experience from latent conceptual representations; hippocampal replay then gradually trains this cortical statistical model [26]. Nagy et al. describe episodic memory as adaptive compression in which surprising information is preserved in relatively raw form whenever cortical models fail to capture important details [1, 27]. A common theme across these accounts is that the statistical regularities stored in cortical weights are leveraged to obtain compressed, distributed memory representations. However, these models credit hippocampus and cortex with complementary but separate representations that are only integrated during retrieval or consolidation.

An important implicit assumption in models that pair feedforward encoding with feedback retrieval is a functional symmetry: the feedback pathway must reconstruct cortical activity from the hippocampal code that was produced by the feedforward pathway. Indeed, O’Reilly and colleagues [28] note that the role of the hippocampal subfield CA1 in inverting the cortical-to-hippocampal mapping remains understudied, despite its centrality to any account of retrieval. A related challenge is computational: if the compression of memories relies on statistical regularities learned through experience, how can the system efficiently represent something it has never encountered before (see also: [1])? We propose that encoding solves these problems through *generative approximation*: rather than relying on a separately trained mapping that directly converts cortical activity into a hippocampal code, the system actively searches for a hippocampal state whose output through a learned cortical generator approximates the current cortical pattern. This analysis-by-synthesis computation [29] allows the system to use prior knowledge to construct representations of novel experiences by flexibly recombining statistical structure already captured in cortical weights.

Generative approximation makes an episodic memory jointly dependent on an episode-specific hippocampal starting point and shared statistical knowledge in cortex. The hippocampal trace identifies a trajectory from which the cortical pattern can be regenerated, while the weights of the cortical generator supply regularities that need not be stored separately in every episodic trace. In this sense, compression is afforded by representing the distinctive content of an episode relative to a generative model of the environment. The precision of the approximation determines the level of detail that can be later recovered, consistent with theoretical accounts positing that memories occupy a continuum from gist-like to highly specific representations [30]. Because encoding searches for a state that will support subsequent generation, it performs a computation closely related to retrieval, and successful encoding can thus be viewed as the first successful attempt to reconstruct the experience. This shared computation is consistent with the generation effect, whereby self-generated information is remembered better than passively studied information [31], and the testing effect, whereby retrieval practice produces greater long-term retention than restudy [32].

Our framework predicts a trade-off between approximation effort and mnemonic precision. Additional search should identify a more accurate generative starting point and support more detailed reconstruction, but at a greater computational cost. This computation-representation tradeoff provides an account for the quality of memory representations that is separable from the quality of their storage [33] and enables an operationalization of “encoding depth”.

We test our predictions in a toy model and in two pretrained generative image models, Stable Diffusion and StyleGAN-XL, which permit stimulus-computable experiments with naturalistic images. First, we show that generative approximation can identify compact starting points for reconstruction without a separately trained feedforward encoder, and that additional approximation improves reconstruction fidelity. In Stable Diffusion, latent representations of semantically related images are well-separated, yet maintain residual category structure. Pronounced category structure re-emerges during reconstruction through the generator, which we take as an analog to memory reinstatement in humans. We then show that reducing approximation produces flatter lure-discrimination gradients in the Mnemonic Similarity Task, mirroring the loss of mnemonic precision observed in aging and mild cognitive impairment. We further find that domain-specific knowledge encoded in the generator weights allows high-quality representations to be approximated more efficiently, capturing the advantage of prior knowledge for memory in a domain of expertise. Finally, we show that representational geometry of latent starting points resembles neural representations in medial-temporal-lobe, and that reinstatement of images through the model traces a reverse visual hierarchy in the brain. Together, these results support an account in which episodic encoding implements a generative search for a compact starting point from which a detailed and distributed cortical representation can later be rebuilt.

### The generative approximation problem to identify episodic codes

We formalized episodic encoding as the problem of finding an episode-specific state that can reconstruct the cortical representation of an ongoing experience through a learned generative model. Let *c* denote the cortical target pattern, *G* a generator whose fixed weights capture statistical regularities learned across experience, and *h* an episode-specific code. The goal of encoding is to identify a state

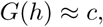

such that passing the stored code through the generator later reinstates the cortical pattern. More formally, the optimal code is

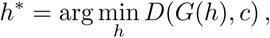

where *D* measures the discrepancy between the generated and target patterns. However, finding the optimal code may require extensive time or computational resources, so the system may instead settle for an approximate code. An approximation procedure *A* produces a code *h_K_* = *A_K_*(*c*; *G*) after *K* computational steps, and retrieval reconstructs the target as *c*^*_K_* = *G*(*h_K_*). The framework therefore postulates that additional approximation should generally improve reconstruction fidelity, and we hypothesize that a generator whose learned statistics match the target domain should require less computation to attain a given level of fidelity. We tested these predictions first in a minimal color model and then in pretrained generative models of natural images.

### Generative approximation identifies reinstatement-capable codes through a fixed generator

To establish the core principle of generative approximation in a minimal setting, we constructed a simple generative network that learned to map a one-dimensional latent code to a structured output. We then used this model to demonstrate how a system can discover compressed representations that enable reconstruction without a dedicated encoder. We envision this as analogous to a neural mechanism of hippocampal generative encoding that exploits cortical knowledge structure rather than relying on an explicit symmetric inverse pathway.

In the notation introduced above, the cortical target pattern *c* was an RGB color, the episode-specific code *h* was an angle on a color wheel, and the generator *G* mapped that angle to its corresponding RGB values (Fig. 1A). Specifically, we instantiated a multilayer perceptron (MLP) that first mapped the latent angle onto its sine and cosine components via a fixed transformation, and then projected these two values through an eight-unit hidden layer onto a target RGB triplet. Importantly, the model was trained only in the generative direction, from angle to color; it was never trained to map an RGB pattern back to its corresponding angle. The model therefore allowed us to test whether a compact, reinstatement-capable code could be discovered through a fixed generator without a separately trained encoder.

**Fig. 1.**
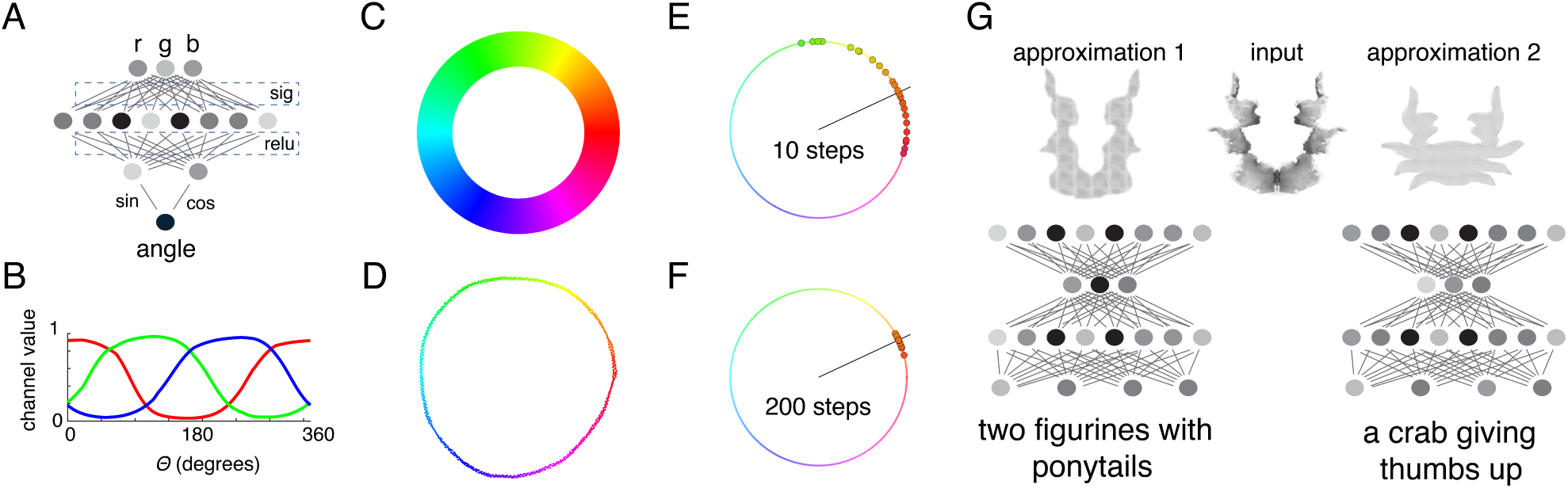
**Toy model and illustration of the generative approximation framework. A,** A simple generative neural network model produces the red, green, and blue components of a color from a latent representation of its angle on the color wheel; the model has only learned a unidirectional transformation from an angle to a color. **B,** The values in the output layer for red, green, and blue channels are displayed for various angles after the model has been trained, demonstrating successful mapping between the latent representation of a color (angle) and its red, green, and blue components. **C,** The HSV color wheel reflects the underlying generative model for the angular representation of color; its structure affords a map for compact latent codes as angles. **D,** t-SNE embedding of the eight hidden-unit activations displays a ring structure that matches the color wheel, demonstrating that the model learns the task structure in its weights. **E, F,** Analysis-by-synthesis encoding of a target color for 25 independent runs. Because the model is only trained in one direction, finding the angle that best represents a color is achieved through iterative gradient-based optimization through the frozen generator weights. The line marks the true angle corresponding to the input color. Points are best approximations for each of the 25 runs from 10 optimization steps (E) and 200 optimization steps (F). More steps yield more accurate latent representations, as evidenced by closer angular distance to the ground truth angle. **G,** Illustration of flexibility in generative approximation. With a larger latent space, different latent representations can yield similar output. For instance, reproducing the never-before-seen input image of an inkblot (here: Card VII of the Rorschach test) via generative approximation could be achieved by a specific instance of two figurines with ponytails, or by a specific instance of a crab giving thumbs up.

The trained generator captured the statistical structure of the color domain. After training on 32,000 samples drawn from the HSV color wheel, the model learned a smooth mapping from latent angle to the red, green, and blue color components (Fig. 1B-C). The network’s hidden representation also reflected this structure: a t-SNE embedding of the eight hidden-unit activation patterns formed a ring consistent with the circular topology of the color wheel (Fig. 1D). Thus, the generator weights encoded the regularities required to reconstruct the three-dimensional RGB color space from a one-dimensional latent variable.

We next fixed the network weights and asked whether encoding could recover the latent angle through generative approximation. Beginning from a randomly initialized angle, the model generated an RGB triplet, computed the reconstruction error relative to the target color, and updated only the angle in the direction that reduced this error. Repeating this analysis-by-synthesis cycle progressively refined the approximate code *h_K_*, bringing *G*(*h_K_*) closer to the target pattern *c*. We compared a condition with 10 approximation steps that yielded a weakly encoded latent (*h*_10_) to a condition with 200 steps that yielded a strongly encoded latent (*h*_200_). Across 25 paired runs, the 200 step condition yielded tight convergence around the target angle (median angular error: 0.86*^◦^*, IQR: 0.18–2.47*^◦^*), whereas 10 steps produced substantially less accurate codes (median angular error: 26.47*^◦^*, IQR: 10.39–37.86*^◦^*). A paired Wilcoxon signed-rank test confirmed that angular error was significantly lower after 200 than after 10 steps (*W* = 325, *p* = 6.54 *×* 10*^−^*^6^, *n* = 25 pairs; Fig. 1E–F). Generative approximation therefore recovered a reinstatement-capable latent code without learning a separate inverse mapping, and additional computation increased the precision of that code.

This simple model instantiates a concrete hippocampal–cortical analogy. The fixed generator weights, trained on the statistics of the color domain, represent slowly learned neocortical knowledge that remains unchanged during the encoding and retrieval of an individual episode [11, 15, 21]. The angle found through approximation represents a rapidly acquired, episode-specific hippocampal trace that serves as a starting point for regeneration. Neither component alone specifies the remembered color: the latent angle identifies the particular color, whereas the generator supplies the shared structure of the color space needed to reconstruct its RGB pattern. The resulting representation is compact because a one-dimensional code can reinstate a three-dimensional target by exploiting statistical regularities captured in the generator weights. Within this architecture, encoding depth reflects how much computation is allocated to identifying the episodic code: deeper encoding yields a more precise starting point for later reinstatement, but requires more processing. Thus, the toy model demonstrates how generative approximation can trade computation for representational precision while distributing a memory across an episode-specific code and a shared generative model.

Although the toy model contains only a one-dimensional latent representation, larger latent spaces provide additional degrees of freedom with which to combine learned statistical knowledge. This flexibility permits multiple latent starting points to yield similar approximations of the same novel input. For instance, the previously unseen inkblot figure on card seven from the Rorschach test [34] could be approximated using a representation resembling two figurines with ponytails, or one resembling a crab giving a thumbs-up (Fig. 1G). In the following section, we explore whether the same computational principle extends to naturalistic images and across generative architectures with substantially richer learned representations.

### Generative approximation extends to naturalistic images across architectures

We next asked whether the generative-approximation principle extends to naturalistic images using deep neural networks capable of generating realistic visual content. We focused on two pretrained image-generation models, Stable Diffusion [35] and StyleGAN-XL [36]. Although these models generate images in different ways, both capture the statistical structure of natural images in their network weights and implement a generative mapping from an image-specific latent state *h* to a detailed image or image-proximal cortical representation *c*. We see the fixed, slowly learned generator *G* as analogous to neocortical knowledge, whereas encoding a particular image requires finding a latent state such that *G*(*h*) *≈ c*. This procedure, known as “image inversion” is typically used for AI-based image editing [37]. In our framework, the resulting latent state serves as the model’s stored memory trace, whereas reconstruction through the generator serves as an analogue of cortical reinstatement.

Stable Diffusion is a text-to-image generative AI model that was trained to reconstruct noise-corrupted images; this reconstruction-training operates on compact latent image-representations derived from a Variational Autoencoder (VAE) that ultimately projects them back into pixel space. During generation, the model begins with a noise-like latent and repeatedly applies the trained denoising network until a clean, image-like latent emerges. Image inversion with Stable Diffusion reverses this generation process. Starting from the VAE-compressed representation of a target image, the model progressively transforms it into a noise-like latent that can later regenerate the target. In the notation introduced above, the clean VAE latent *z*_0_ serves as the model’s cortical target *c*, whereas the noise-like inversion endpoint *z_T_* serves as the episode-specific starting point *h*; after reconstruction, the VAE decoder maps the recovered *z*^_0_ back into pixel space [35, 38].

StyleGAN-XL implements the same computational objectives through a different mechanism. The model generates an image by passing a latent code through a sequence of style-modulated layers. Earlier layers primarily determine coarse properties such as spatial layout and object shape, whereas later layers add progressively finer features such as texture and local detail.

To implement inversion of a target image in StyleGAN-XL, we began with an initial latent code, generated an image from it, and compared the generated image with the target. The latent code was then iteratively adjusted to reduce the discrepancy while the generator weights remained fixed [39], much like the angle-to-RGB approximation in Fig. 1. StyleGAN-XL can therefore directly instantiate analysis-by-synthesis: each update evaluates how well generation from the current code approximates the observed image. Stable Diffusion instead obtains a starting point by traversing a structured inverse diffusion trajectory. Despite this algorithmic difference, both methods implement the same computational goal of finding an *h* for which *G*(*h*) *≈ c*.

We define encoding depth as the number of approximation steps applied during image inversion; we envision this as the computational analogue of encoding depth in biological memory. Deep encoding produces latent representations that yield a high reconstruction fidelity while shallow encoding produces latent representations that lead to impoverished reconstruction (Fig. 2A–B). We further envision the reconstruction process of passing the latent through the weights of the network as a computational analogue of memory reinstatement.

**Fig. 2.**
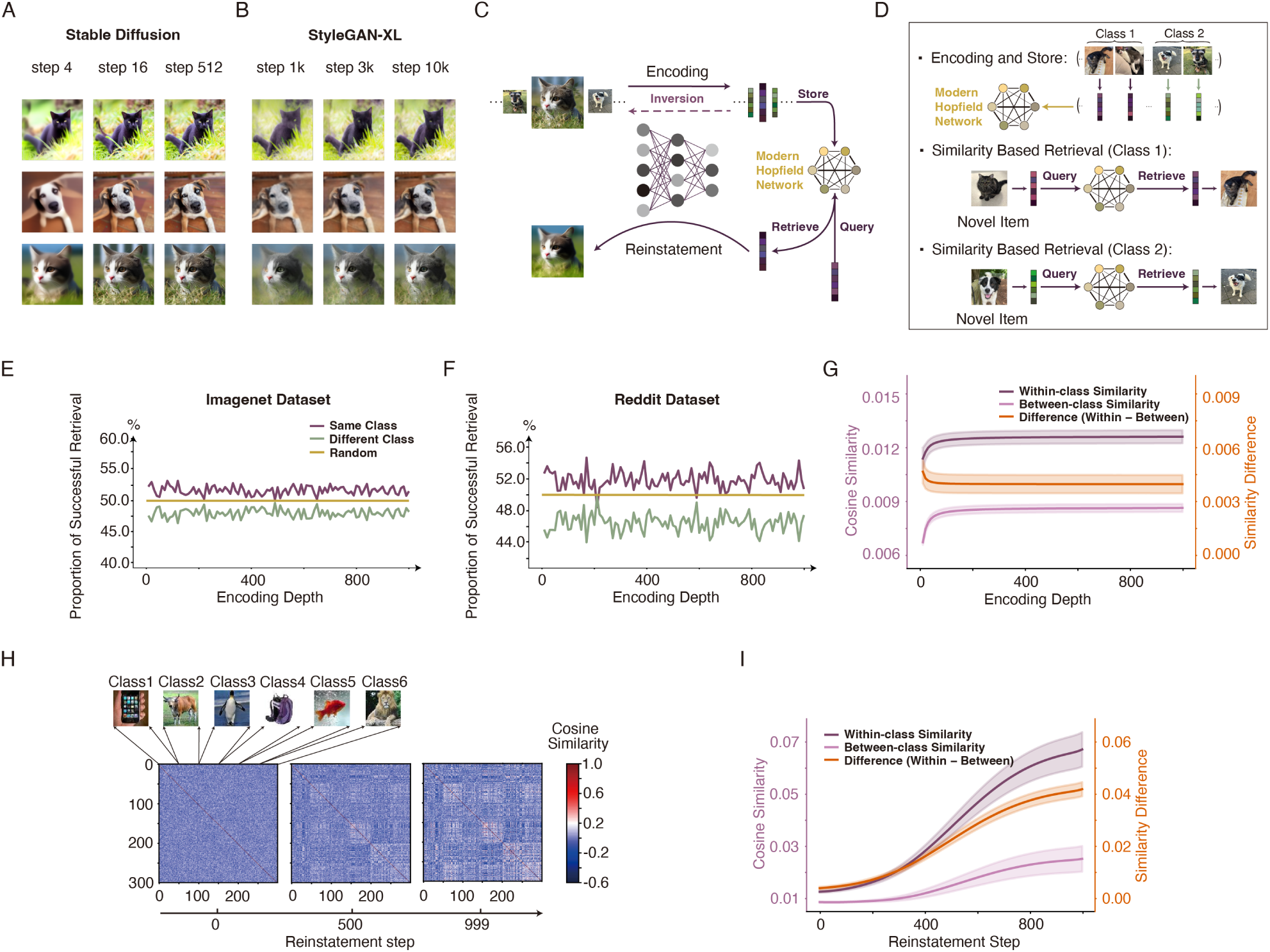
**Generative approximation produces well-separated codes with minimal category structure that is reexpressed during reconstruction. A,** Example reconstructions of three natural images using Stable Diffusion with increasing numbers of DDIM inversion steps (shown: 4, 16, and 512 steps). Deeper inversion yields progressively more faithful reconstructions. **B,** Corresponding reconstructions using StyleGAN-XL after 1,000, 3,000, and 10,000 latent-optimization steps. Increasing approximation improves the quality of the recovered image. **C,** Schematic of our complementary memory architecture. A target image is encoded by inversion into an episode-specific latent starting point, the latent is stored in a Modern Hopfield Network, and retrieval reinstates the image by passing the recovered latent through the same generative model. **D,** Similarity-based list-learning paradigm. Two images from different classes are encoded and stored in the MHN. A novel image from one of the classes serves as a query to test whether the corresponding class representation is preferentially retrieved from memory. **E,** Retrieval performance (percentage of all trials) as a function of diffusion depth for in-domain Tiny ImageNet images and out-of-sample Reddit images (**F**). A same-class preference persists across inversion depths, indicating preservation of category structure in the stored latent code. **G,** Cosine similarity between images within the same class remains above between-class cosine similarity across all levels of encoding depth; their difference remains stable after the first 50 inversion steps. **H,** Time-resolved representational similarity matrices for diffusion latents throughout the reinstatement process from the deepest inversion latent (999 step inversion). Category block structure re-emerges during reinstatement. Matrices are shown for one of the five image sets at reconstruction steps 0, 500, and 999 (from left to right). **I,** Mean within-class and between-class similarity across reinstatement; shaded bands show the standard error across the five image sets. The increase in within-class similarity exceeds the increase in between-class similarity, which quantifies how semantic category structure is re-expressed throughout the image reconstruction process (a computational analogue of cortical ”reinstatement”).

### Similarity based retrieval from reinstatement-capable codes

We next implemented a full model where latent starting points found through image inversion are embedded within a complementary memory architecture and function as hippocampal memory traces. In the full model, the weights of the generative model served as slowly learned neocortical knowledge, while a Modern Hopfield Network (MHN) stored the image-specific latent states instantiating fast hippocampal storage (Fig. 2C). Because of the computational demands of the subsequent simulations, we used Stable Diffusion as the cortical generator throughout these analyses. At retrieval, a cue was used to select a stored latent from the MHN, and the recovered latent was then passed through the same generative model that had been used to compute it, reinstating the corresponding image representation. This architecture therefore separates rapid storage of episode-specific starting points from the shared statistical knowledge used to reconstruct their content.

We first asked whether similarity among the stored latent states remained sufficient to support category-guided retrieval. In a list-learning experiment, two images from different categories – for example, a cat and a dog – were encoded through image inversion and their latent states were stored in the MHN. Retrieval was then probed using the latent state of a novel image belonging to one of the two studied categories (Fig. 2D). We scored retrieval as category-consistent when the probe was more similar to the stored representation from the same category than to the representation from the other category. We performed this analysis using both in-domain images from Tiny ImageNet and novel out-of-sample images collected from Reddit after the model’s training-data cutoff. Probing with a different exemplar, rather than the exact studied image, allowed the experiment to measure similarity in the latent space instead of item-matching with trivially perfect performance.

Shallow inversion states retained sufficient semantic structure to preferentially retrieve memories from the same category: Across inversion stages, the proportion of same-category retrieval was 51.8% for Tiny ImageNet and 52.7% for the Reddit images, both significantly above the chance level of 50% (one-sided binomial, *p <* 0.001), and did not change across encoding depths. This was evidenced by near identical performance in the first and last third of encoding depths (51.9% and 52.7% for the Tiny ImageNet and Reddit images, respectively; Fig. 2E-F). Additionally, the correlation between encoding depth and same-class retrieval was *r*= *−*0.05 for Tiny ImageNet and *r* = *−*0.00 for the Reddit images, supporting the notion of preserved performance (*BF*_01_ = 7.2, and 8.0, respectively). Examining the similarity between images directly, we found persistently higher similarity between within-class images than between images from different classes at every encoding depth (one-sided paired *t*-tests across the five image sets, all *q <* 0.002). After a small decrease within the first 50 encoding steps, the within-minus-between difference did not change further with increasing encoding depth (fitted linear change of *−*0.9% of the mean difference across encoding depths 50 to 999; statistically equivalent to zero within bounds of *±*10%, two one-sided tests across the five image sets, both *p <* 0.001; Fig. 2G). These findings demonstrate that latent codes can retain the category structure that is needed for similarity based retrieval from semantically related cues [40]. Importantly, category structure re-emerged when the stored latent states were passed forward through the generative model. During the reconstruction of latents from the deepest inversion state, both within-category and between-category similarities increased (both Spearman’s *ρ ≥* 0.998, *p <* 10*^−^*^10^), and within-category similarity diverged from between-category similarity (Fig. 2H-I). The within-minus-between gap widened with a slope of *β* = +4.51 *×* 10*^−^*^5^ per reconstruction step, and the standardized difference reached Cohen’s *d* = 6.92 by the end of reconstruction. Thus, although latent states did not themselves express a strongly category-organized geometry, they retained sufficient episode-specific information to recover that structure when combined with the statistical knowledge stored in the generator weights.

Together, these findings describe a memory architecture, in which generative approximation produces well-separated and item-specific starting points that are stored in hippocampus. These latent starting points preserve enough semantic structure to allow for retrieval from semantically similar cues. Shared semantic structure is then re-expressed during cortical reinstatement.

### Depth of encoding is a computational account of mnemonic precision

We next asked whether the depth of generative approximation could serve as a computational analogue of encoding strength. In episodic memory, stronger encoding should preserve the fine-grained details needed to distinguish an old item from a highly similar lure, whereas weaker encoding should result in a more gist-like representation [2, 3]. In our framework, encoding depth corresponds to the amount of computation devoted to finding an episode-specific code *h_K_* for which *G*(*h_K_*) approximates the target representation *c*. We operationalized encoding depth as the number of approximation steps *K*: deeper encoding should yield a more precise starting point for reconstruction, whereas shallower encoding should yield a less precise code. We tested this prediction using the Mnemonic Similarity Task (MST), which measures mnemonic precision by asking participants to discriminate previously studied objects from perceptually similar lure items [41].

The MST provides a graded test of how much item-specific information is retained in memory. Lure items are grouped into five similarity levels: Lure 1 items are most similar to their studied counterparts, whereas Lure 5 items are least similar. Repeated items are therefore most likely to be correctly identified as old, highly similar lures are more likely to be falsely endorsed as old than less similar lures, and novel items are least likely to be endorsed. Older adults and patients with mild cognitive impairment (MCI) typically exhibit a flatter, sublinear gradient across these conditions, reflecting disproportionate difficulty rejecting highly similar lures [42] (Fig. 3C). We therefore asked whether reducing encoding depth in the generative-approximation model was sufficient to produce a similar selective loss of mnemonic precision.

**Fig. 3.**
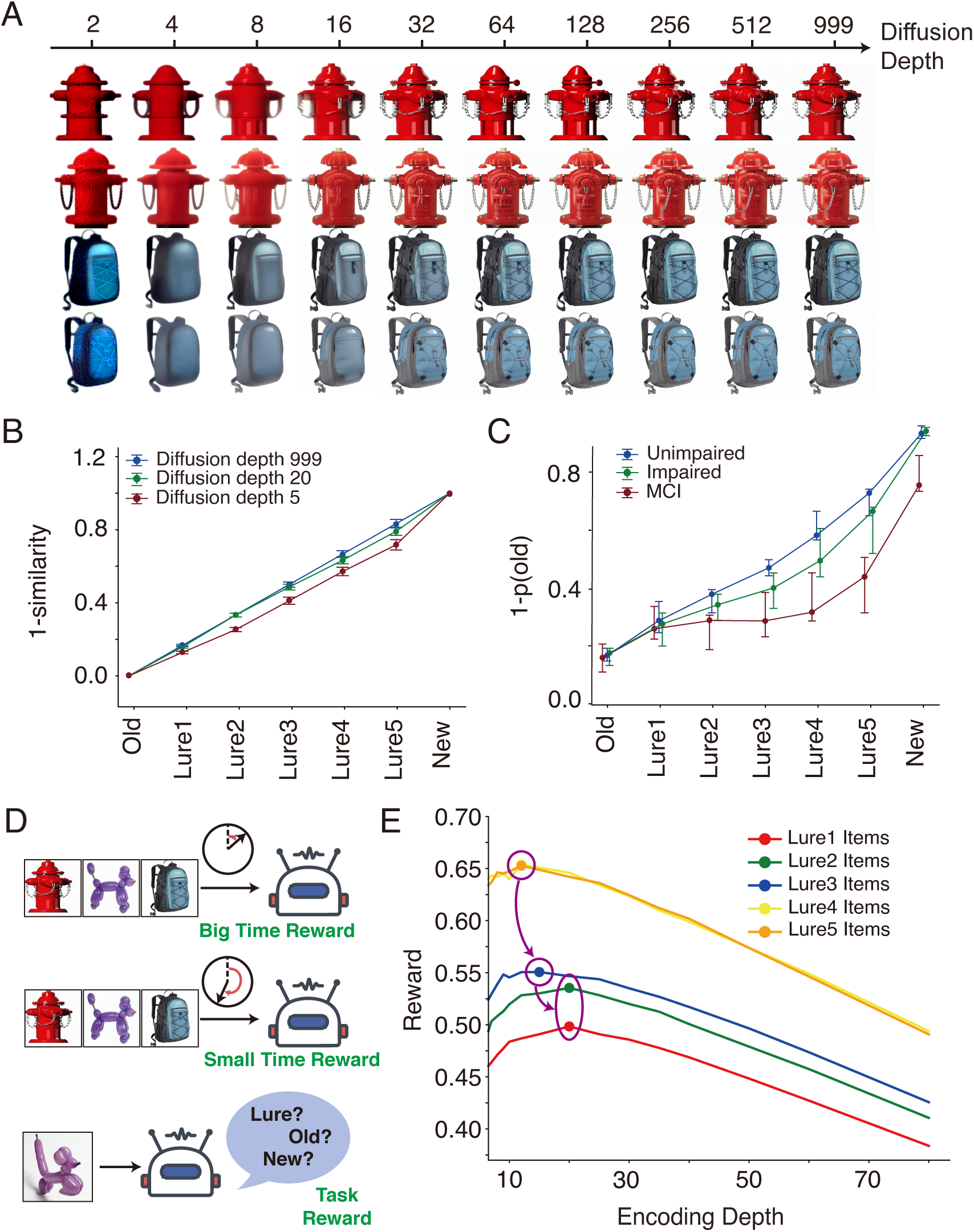
**Encoding depth determines mnemonic precision. A,** Example reconstructions for two MST old-lure pairs across encoding depths. Shallow encoding preserves primarily coarse shared structure, whereas deeper encoding progressively recovers the item-specific details needed to distinguish studied objects from highly similar lures. **B,** Model-based dissimilarity (1 *− s*^) between images of the same object for deep, intermediate, and shallow encoding conditions (999, 20, and 5 diffusion steps), with repeated items anchored at 0 and novel foils at 1 so that the orientation matches C. Probe–memory dissimilarity increases from repeated (old) items through increasingly distinct lures to new foils, and this gradient becomes progressively flatter as encoding depth decreases. Lure levels are rescaled per level to a linear increase in the deepest condition to facilitate comparison with C; error bars show the standard error across test probes. **C,** Graphical illustration of empirical MST performance curves for unimpaired, impaired, and MCI groups, shown in the same ordered format (illustrating findings from: [41]). The flatter profile associated with reduced mnemonic precision qualitatively parallels the effect of shallow encoding in the model. **D,** Schematic of the encoding-depth tradeoff between representation and computation. Shallower encoding is associated with a larger time reward because it is computationally cheaper, whereas deeper encoding yields more precise memory traces but a smaller efficiency reward; total task reward is obtained by combining the reward for time and accuracy.**E,** Total reward as a function of encoding depth for lure levels 1–5. If total reward combines lure-discrimination performance with a decreasing time-reward schedule that imposes a penalty on computation, the reward-maximizing depth shifts systematically deeper as lure similarity increases.

We first verified that encoding depth controlled reconstruction fidelity for the MST stimuli. We used 999, 20, and 5 DDIM inversion steps to define deep, intermediate, and shallow encoding, respectively. Each studied image was inverted to obtain an episode-specific code *h_K_* and then reconstructed as *ĉ_K_* = *G*(*h_K_*) (Fig. 3A). As encoding depth *K* increased, the model progressively recovered the diagnostic visual details of the studied objects. Across 256 study images, reconstruction fidelity increased with encoding depth, and deep encoding produced markedly more faithful reconstructions than shallow encoding (999 versus 5 steps; mean Structural Similarity Index Measure, SSIM: 0.86 versus 0.69; paired Wilcoxon signed-rank test, *p <* 0.001; [43]). Thus, increasing encoding depth improved the fidelity of the representations available at retrieval.

We next tested whether encoding depth altered the representational evidence available for mnemonic discrimination, when performing the MST [42]. During the encoding phase, the final inversion latent *h_K_* for each studied image served as its memory trace; at test, the trace was reinstated through the diffusion model, yielding *ĉ_K_*, and each probe was compared with the reinstated trace of its own studied item. We computed the cosine similarity between the reconstructed memory *ĉ_K_* and the initial VAE representation *z*_0_ of each test probe, which could be an exact repetition (old), one of five levels of similar lure (L1–L5), or a novel item (new). At every encoding depth, reconstructed memories were most similar to repeated items, showed intermediate similarity to lure items, and were least similar to novel items (Welch’s *t*-tests on raw cosine similarity with one value per image; old vs. lures, *t ≥* 35.9; lures vs. new, *t ≥* 20.6; all *p <* 10*^−^*^85^). Thus, the model produced the basic ordering of mnemonic evidence required to distinguish repeated items, similar lures, and novel foils.

We then expressed probe–memory similarity relative to the mean similarities of old (1) and new items (0), placing the model’s responses on a common recognition-strength continuum analogous to the signal-detection problem posed by the behavioral MST [42, 44]. Mean anchored probe–memory similarity across Lure 1–Lure 5 was 0.420, 0.376, 0.372, 0.263 and 0.271 after 999 inversion steps; 0.450, 0.377, 0.390, 0.300 and 0.308 after 20 steps; and 0.555, 0.526, 0.484, 0.368 and 0.372 after 5 steps. That is, reducing encoding depth brought lure levels closer to the old item (repeated-measures ANOVAs, *F* (2, 296–322) = 63.6–141.1, Greenhouse–Geisser corrected, all *p <* 10*^−^*^19^, 149–162 pairs per level); pairwise contrasts confirmed this ordering at every lure level (paired *t*-tests; 20 vs. 999 steps, *t ≥* 2.77, all *p <* 0.01; 5 vs. 999, *t ≥* 10.05; 5 vs. 20, *t ≥* 8.06; both *p <* 10*^−^*^12^).

Because repeated and novel items are fixed by the anchoring, this displacement of the intermediate lure levels is precisely a flattening of the lure-discrimination gradient (Fig. 3B, note that the lines in B have been rescaled per lure level to a linear increase in the deepest condition to facilitate comparison with C). Shallow encoding therefore impaired the representational precision needed to distinguish highly similar lures from studied items. Indeed, the loss of mnemonic precision produced by shallow encoding resembled the flattened MST profiles observed in aging and MCI (Fig. 3C). In both the model and the empirical profiles, the discrimination between studied and novel items remained possible, whereas similar lures became selectively more difficult to distinguish from stored memories [41, 42].

Having established that deeper encoding improves mnemonic precision, we next demonstrate how a resource-limited system should allocate encoding depth. Deeper encoding produces more faithful memory representations, but it also requires additional computation. We formalized this trade-off by combining a discrimination reward, computed as the complement of normalized lure similarity, with an efficiency reward that decreased monotonically with encoding depth (Fig. 3D). The combined objective therefore captured the tension between representation quality and computational cost: deeper encoding improved discrimination, but produced diminishing total returns as its computational penalty increased. Note that the goal of this analysis is to illustrate how a cost of computation could shape encoding policy, rather than to estimate the brain’s actual cost function. Through this time-dependent reward weighting, easy lures can achieve their highest total reward at relatively shallow encoding depths, whereas increasingly confusable lures justify progressively deeper encoding (Fig. 3E).

In our example, the reward-maximizing depth becomes ordered by lure difficulty: Lure 1 and Lure 2 attained their maximum total reward after 20 encoding steps, Lure 3 after 15, and Lure 4 and Lure 5 after 12 (Spearman’s *ρ* = *−*0.95 between lure level and reward-maximizing depth). Because the reward function is evaluated analytically over a stipulated cost schedule, we only report this ordering descriptively. This analysis illustrates that, once computation carries a cost, deeper encoding is not uniformly optimal. We argue that memories should be encoded only as deeply as required to support the precision expected to be useful at retrieval.

### Memory representations become functionally distributed across the latent trace and generator weights

To understand how the episode-specific latent code and the shared generative model jointly support memory reconstruction, we next characterized the transformation of image specific information across the network. In the generative-approximation framework, the latent code *h_K_* contains information that distinguishes a particular episode, whereas the fixed generator *G* contains statistical knowledge learned across many experiences. The reconstructed representation *ĉ_K_* = *G*(*h_K_*) therefore depends on both components: *h_K_* specifies which episode should be reconstructed, while *G*(.) transforms that code into a structured, image-like representation. In this functional sense, a memory representation is distributed across an episode-specific code and shared knowledge in the generator, i.e., while all image-specific information is entailed in *h_K_*, it needs to be decoded through the nonlinear transformations of the network.

To characterize this process, we estimated two mutual-information quantities across encoding depths. Let *c* be defined by the initial image-proximal VAE embedding *z*_0_ in Stable Diffusion, with *h_K_* denoting the inversion latent obtained after *K* encoding steps, and *ĉ_K_* = *G*(*h_K_*) denoting the representation recovered after reconstruction. Using InfoNCE, which provides a lower-bound estimate of mutual information in high-dimensional spaces, we computed two separate measures:

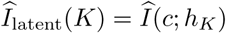

and

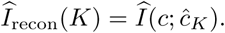

The first measure quantified how detectable image-specific information remained in the episode-specific latent, whereas the second quantified how detectable that information became after the complete encoding– reconstruction cycle (Fig. 4A) *Î_l_*_atent_(*K*) declined monotonically with encoding depth (*p <* 0.001); conversely, *Î*_recon_(*K*) increased with encoding depth (*p <* 0.001, Fig. 4B). Thus, deeper encoding produced latent states in which image identity was less directly detectable, while simultaneously producing reconstructions in which images could more reliably be identified. Note that higher detectable information in the reconstructed image compared to the latent is not at odds with the data processing inequality, which states that processing a signal cannot increase the amount of information it contains [45]; information is still preserved in the latent *h_K_*, however, recovering the image requires a multi-step non-linear decoding operation, which is realized by the weights of the network [46]. Importantly, the crossover between the trajectories of *Î*_latent_ and *Î*_recon_ marks the encoding depth at which regeneration through *G*(.) improves accessibility of image content over the latent trace. We defined the *crossover depth*, *K*_cross_, as the encoding depth at which *Î*_recon_(*K*) first equaled or exceeded *Î*_latent_(*K*). *K*_cross_ provides an index of the division of labor in a distributed memory system; it captures the computational investment required before shared statistical knowledge, rather than the memory trace itself, carries the burden of making an episodic memory’s identity accessible.

**Fig. 4.**
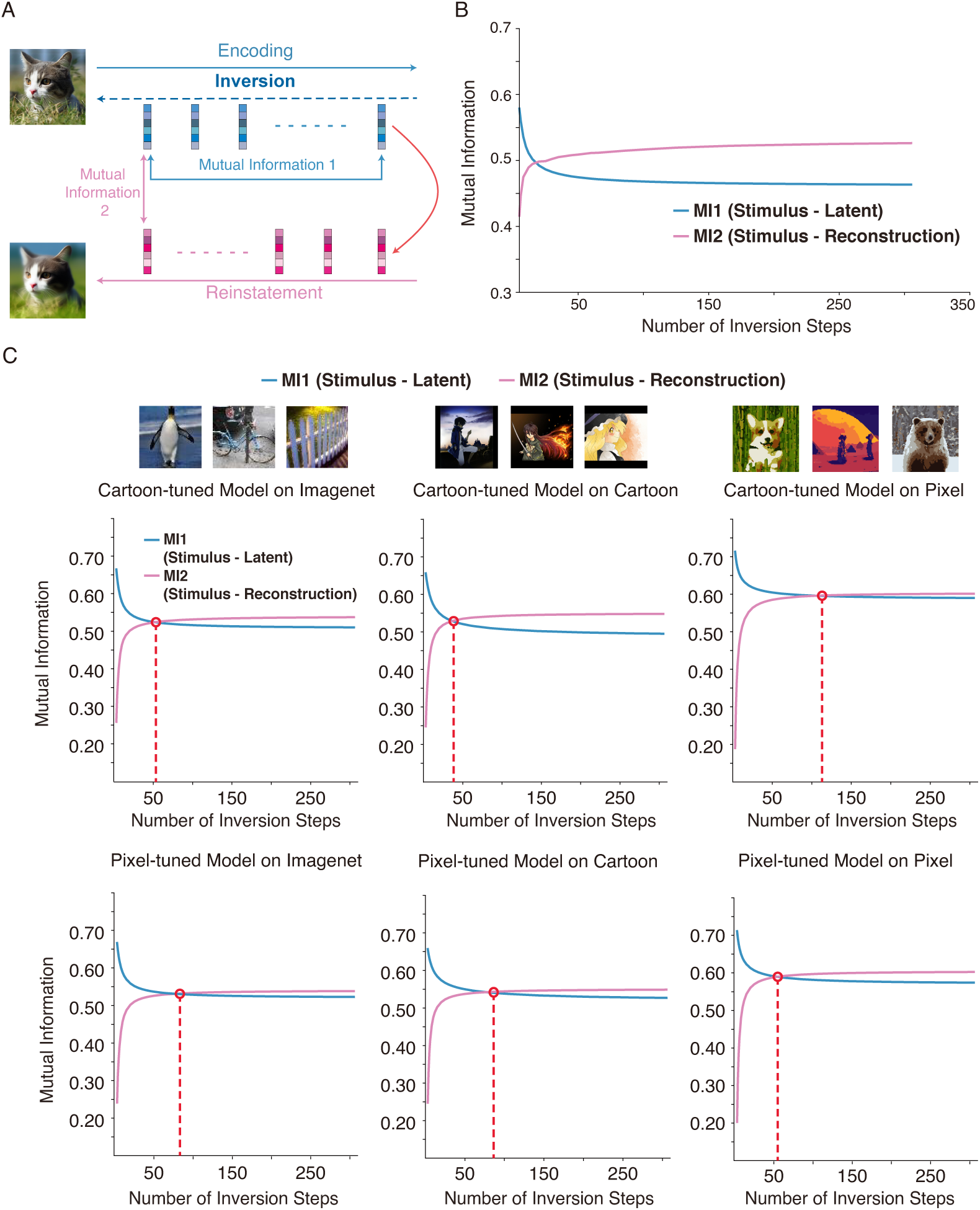
**Domain-matched generative knowledge supports reconstruction at shallower encoding depths. A,** Schematic of the two InfoNCE-based mutual-information estimates used to characterize the encoding–reconstruction cycle. An image-proximal representation *c* is inverted into an episode-specific latent *hK* and subsequently reconstructed as *ĉK* = *G*(*hK*). *Î*latent estimates how directly image identity can be detected from the inversion latent, whereas *Î*recon estimates how readily it can be detected after reconstruction. Their crossover marks the encoding depth at which the reconstruction estimate first equals or exceeds the latent estimate. **B,** Information estimates across encoding depths for the untuned Stable Diffusion model on naturalistic images. **C,** Information estimates across tuned-model and stimulus domains. Rows correspond to the models (cartoon-tuned model and pixel-tuned model), and columns correspond to image domains (natural images, cartoon images, and pixel-art images). Dashed red lines and circles indicate crossover depths. The cartoon-tuned model crosses earliest for cartoon images, whereas the pixel-tuned model crosses earliest for pixel-art images. Thus, each model reaches its crossover depth soonest in its domain of expertise, demonstrating an expert effect in generative memory encoding.

### Domain-matched expertise of the generator supports efficient computation of distributed representations

Based on empirical evidence from human psychology research [47–50], we hypothesized that the computational investment required to reach the crossover point – and consequently to learn a distributed representation – would depend on the domain of expert knowledge that the generator has stored in its weights.

We therefore examined whether *K*_cross_ depended on how well the generator’s existing knowledge matched the image domain. To this end we fine-tuned Stable Diffusion separately on cartoon and pixel-art images using Low-Rank Adaptation adapters [51]. This procedure yielded two domain-specific expert models whose generator weights were adapted to cartoon and pixel-art statistics, respectively. We crossed the untuned, cartoon-tuned, and pixel-tuned models with natural (ImageNet), cartoon, and pixel-art stimuli, producing nine model–stimulus conditions. Because adapters are trained with domain-specific caption tokens, images in this experiment needed to be encoded with a short domain prompt – unlike the unconditional setting used before. Critically, the prompt was tied to the stimulus domain, not to the model, i.e., for any given image set, all three models received the identical prompt, so differences in crossover depth across models can only reflect their adapted weights.

In the untuned Stable Diffusion model, the crossover *K*_cross_ occurred on average after 18.2 (SD = 3.7) encoding steps across five replicates for natural images from the model’s native training distribution (Fig. 4B), after 28.2 (SD = 3.8) steps for stylized cartoon images, and after 29.9 (SD = 5.4) steps for pixel-art images. The crossover was thus earliest for the image domain that dominated the model’s pretraining corpus. The domain-matched generators reached *K*_cross_ at earlier encoding depths than mismatched generators. The cartoon-tuned model’s *K*_cross_ was earliest for cartoon images, after 38.5 (SD = 8.2) encoding steps, compared with 53.3 (SD = 9.5) steps for natural images and 113.1 (SD = 31.4) steps for pixel art. Similarly, the pixel-tuned model’s *K*_cross_ was earliest for pixel-art images, after 54.9 (SD = 6.8) steps, compared with 83.1 (SD = 29.8) steps for natural images and 86.4 (SD = 36.1) steps for cartoons (Fig. 4C). To evaluate this matched-versus-mismatched effect statistically we subtracted the summed *K*_cross_ of the mismatched conditions (cartoon-on-pixel plus pixel- on-cartoon) from the matched conditions (cartoon-on-cartoon plus pixel-on-pixel). The observed difference of *−*106.17 steps indicates that in matched conditions *K*_cross_ occurs earlier than in mismatched ones. We then generated a noise distribution of this contrast by shuffling the model-to-image-domain association 10,000 times and computing surrogate differences. The observed value was significantly more extreme than the values from the random distribution (*p <* 0.001), indicating that each network encoded the stimuli most quickly for the domain whose statistics it had learned best.

These results show how statistical knowledge stored in a generator can function as a computational resource for episodic encoding. When the generator’s learned statistical weights were well-matched to the stimulus domain, the model required fewer processing steps to leverage statistical knowledge effectively to represent an episodic memory.

### Generative representations have MTL-like geometry, their reinstatement retraces a reverse visual hierarchy

We finally asked whether the latent representations learned through generative approximation resemble those observed in the human brain. To test this, we conducted a systematic comparison to functional magnetic resonance imaging (fMRI) data from the Natural Scenes Dataset (NSD; [52]). The NSD contains large-scale 7-Tesla fMRI recordings of eight participants who performed a continuous recognition task with thousands of natural images; participants judged whether images were new or had been presented at any previous point in the experiment. We computed latent representations for all images shown in the analyzed sessions and used a cross-modal representational analysis to compare them to the neural recordings [53]. We asked how the representational geometry of the deepest latent compared to neural activity patterns for various regions of interest. In a second analysis, we compared the representational geometries between neural activity and the intermediate latent representations throughout the reconstruction process from the deepest latent representation. To this end, we extracted the Stable Diffusion latents at the deepest encoding depth (999) and from 34 evenly-spaced states (every 30 steps) along the reconstruction trajectory (starting from the deepest inversion state); we then computed the trial-by-trial representational similarity matrices across images from the model’s latent representations. For comparison, we computed the trial-by-trial similarity matrices derived from multivoxel response patterns in anatomically defined visual and medial temporal regions of interest (see Methods). Model–brain correspondence was quantified with centered kernel alignment (CKA, [54]), which measures the similarity between two representational geometries while remaining invariant to rotation and isotropic scaling, making it suitable for comparing spaces with different dimensionalities and coordinate systems (Fig. 5A). This analysis therefore tested whether relationships among remembered images change across the generative trajectory in a manner that parallels relationships expressed across the visual-medial temporal hierarchy.

**Fig. 5.**
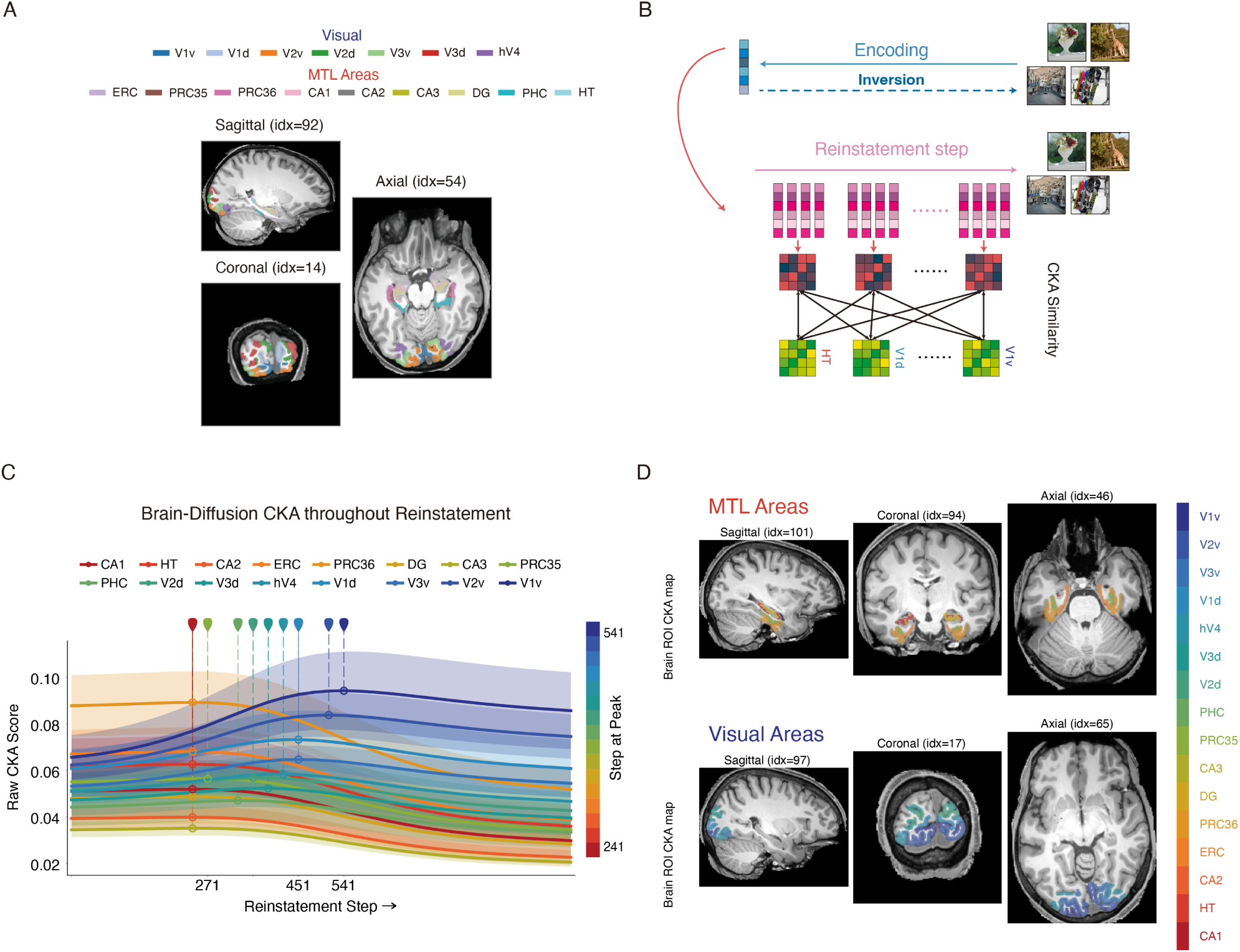
**Generative approximation produces MTL-like codes; reconstruction traverses the visual hierarchy in reverse order. A,B** Schematic of model-brain representational similarity analysis. The same set of NSD images was passed through diffusion inversion and reinstatement, and latent states were sampled across the reconstruction trajectory. At each sampled depth, a trial-by-trial representational similarity matrix was computed from the model latents. For the same image trials, multivoxel activity patterns were extracted from anatomically defined visual and medial temporal ROIs (**A**), and corresponding neural similarity matrices were computed. Model and brain representational geometries were then compared using centered kernel alignment (CKA; **B**). **C,** Model– brain correspondence during reinstatement. Raw CKA scores are sampled across reconstruction steps. The ordering of peak-alignment follows a reverse visual hierarchy, consistent with a return from well-separated memory-like representations toward more visual cortical formats during reconstruction. Curves show the mean across the eight participants and shaded bands *±* 1 s.d.; peak markers show the peak of the group-average CKA curve for each ROI, and colors rank the ROIs by this peak (dark red: earliest, dark blue: latest). **D**: anatomical maps of reinstatement peak depth, colors indicate the rank of the reconstruction step at which each region has its maximal CKA alignment score (colors as in C).

Our central prediction was that peak model–brain correspondence of the latent starting point would fall into the medial temporal lobe, where neural codes are well-separated into item specific codes [55]. Throughout the reinstatement process, we predicted that the format of the latent representation would trace a reverse visual hierarchy. Consistent with this prediction, the latent starting point had the highest representational similarity with the perirhinal cortex (PRC36; highest in seven of eight participants), and throughout the reconstruction steps CKA between the model’s latent representations and the neural data peaked at different reconstruction steps for different regions (Fig. 5C).

We first verified that alignment reflected image-specific correspondence rather than incidental similarities by shuffling which fMRI response was paired with which model representation and recomputing CKA on 1,000 permutations. Similarity between neural representations and model representations exceeded chance level throughout (one-sided permutation tests, all *p ≤* 0.022, significant after false-discovery-rate correction at *q* = 0.05).

To quantify whether the ordering of peak alignments followed the reverse cortical processing hierarchy, we assigned each region a rank according to its position in the canonical visual hierarchy (see Methods), with subdivisions at the same hierarchical level sharing a rank [56]. Indeed, the ordering of peak correspondences reversed along the cortical trajectory (Fig. 5C): MTL regions peaked earlier during reconstruction, whereas visual regions peaked later as the representation approached the sensory format. Cortical hierarchy rank was strongly negatively correlated with reconstruction peak depth (Spearman’s *ρ* = *−*0.90, permutation *p <* 0.001). V1 peaked at a mean reconstruction step of 485, whereas MTL regions peaked at step 252 (paired *t* = 20.88, *p* = 1.5 *×* 10*^−^*^7^; Wilcoxon signed-rank test, *W* = 0, *p* = 0.008). Thus, reconstruction progressed from an MTL-like toward an increasingly visual-like representational geometry. While we do not wish to imply that the brain implements the specific DDIM algorithm or that any individual diffusion state is identical to a hippocampal representation, these findings provide neural support for the proposed representational organization.

## Discussion

We model the encoding of episodic memory representations as an approximation problem in generative neural network models. The resulting memories that are stored in a model of hippocampus are neither exact copies of sensory input, nor indices that point to cortical patterns. Instead, we find compact “starting points” that can reproduce activation patterns in a neocortex, which is implemented by the generative model. In a minimal color generation model and in large image generation AI models, we demonstrate how these starting points can be learned on a trial-by-trial basis by optimizing for the ability of the network to re-generate cortical patterns. This *generative approximation* framework unifies encoding and retrieval as a single generative operation.

A central theoretical contribution of our work is that it describes an account of memory encoding without a feedforward encoder. In computational models of episodic memory, encoding typically proceeds by transforming sensory input through a cortical-to-hippocampal pathway and retrieval inverts this transformation via back-projections [11, 22]. An implicit assumption of these architectures is symmetry between two learned mappings: bidirectional mappings must remain mutually invertible. By discovering representations through generation rather than through symmetric inversion our framework dissolves this problem, achieving flexibility in mnemonic precision in a single system by replacing a deterministic invertible mapping with approximation [29, 37, 39, 57]. Generative approximation consequently formalizes a tradeoff between computation and mnemonic precision: episodic memory encoding becomes a costly search process to derive more accurate and detailed memory representations. We modelled seminal findings from the mnemonic similarity task (MST) with this approach: lower encoding depths increased similarity of similar lure items, moving them closer to identical repeats on the similarity-continuum between old and new items. This modeling result mirrors behavioral findings from aging adults and patients with mild cognitive impairment, who selectively struggle with the discrimination of similar lure items from previously studied (old) items [41, 42]. Our findings explain MST deficits with a reduction in encoding depth that shifts the detail of memories toward gist-like representations, consistent with eye-tracking evidence that failed lure discrimination reflects impoverished encoding [58]. This correspondence does not imply that reduced encoding depth is the sole mechanism underlying age-or disease-related memory impairment; rather, it demonstrates that a reduction in encoding computation alone can shift the representational format of memory toward a more gist-like regime without requiring a qualitatively different retrieval operation (see also [59]). Thus, generative approximation provides a computational mechanism by which shallow encoding can produce a qualitative signature of diminished mnemonic precision. We further demonstrated how this tradeoff can be embedded in a resource rational framework [60]. If computations are allocated optimally based on temporal constraints or memory demands, detailed learning of memory representations may only be performed to the extent it is needed at retrieval (see also: [61]).

Our analyses further demonstrate how generative encoding leverages the division of labor across the components of the memory system. With deeper encoding, the detectable image-information in the hippocampal latent decreased, yet sufficient residual category structure remained to allow for retrieval based on semantic similarity, which is in line with recent evidence that neural populations in hippocampus retain semantic structure [40]. Generative encoding consequently produces latent representations that are transmuted from copy-like embeddings to well-separated latent codes. The amount of detectable information recovered through reconstruction, on the other hand, increased with encoding depth, indicating that the cortical generator assumes a progressively larger role in the reinstatement of memories. Domain expertise facilitated this division of labor. Specifically, when the learned statistics of the generator matched the stimulus domain of the memory, the point of approximation where the cortical generator assumes an important role in reconstructing image identity was reached with less computational investment. This finding provides a computational account of the well-documented phenomenon that experts form precise domain-relevant memories more efficiently than novices [47–50], and is consistent with evidence that schema-congruent information is encoded more rapidly when relevant cortical knowledge structures are available [62].

Our simulations largely rely on generative AI models that were not designed as neural process models. Moreover, trajectory-based inversion (DDIM inversion) [57] is not itself a plausible neural process, which is why we further demonstrated the framework with a toy model and optimization-based inversion in StyleGAN-XL, which improves precision by iterating rather than by adding network depth. Note, however, that optimization-based inversion can also be used to compute Stable Diffusion image inversion [63] but we used DDIM for computational efficiency. Crucially, the representational alignment between Stable Diffusion and human 7-Tesla fMRI data provides evidence that our framework captures a shared structure with neural codes for memories in humans. Specifically, the representational geometry of the model’s latent starting point corresponded to regions of interest in the medial temporal lobe. During reconstruction, the representational format of memory reconstruction in the model then tracked the cortical hierarchy in reverse order: medial temporal regions aligned with early reconstruction stages and visual areas aligned optimally with late stages, tracing the trajectory from an MTL-like code toward sensory-like representations. This bidirectional pattern situates the model’s latent at the apex of a processing hierarchy that mirrors the anatomical organization between occipital cortex and hippocampus, and characterizes cortical reinstatement as a plausible neural process for the reconstruction of detailed memories from compact starting points. These findings converge with recent brain decoding work demonstrating that diffusion model latent spaces can reconstruct perceived images from fMRI activity [64, 65], which provides independent evidence that model representations in generative AI models occupy a neurally meaningful space (see also: [66]). A key characteristic of our modelling approach is that we use the same set of cortical weights to store all episodic memories, whereby the content of episodic memories becomes jointly represented by the latent starting point and the weights of the cortical generator. In this architecture a change of cortical weights and a change in the latent “starting point” will both distort the content of existing episodic memories, unless these changes are offset by corresponding update processes, such as offline replay [67]. Nevertheless, a mismatch in hippocampocortical correspondence may regularly occur in the brain. For instance, we often experience surprise, when an item that we remember from childhood (e.g., a chair) suddenly appears much smaller than we remember it; a gradual change in cortical representations of objects may not have been reconciled with the hippocampal memory trace.

A complementary framing for our findings is the idea of transfer appropriate processing, which describes that it is beneficial to learn in a way that resembles later retrieval [68, 69]; our generative approximation framework implements a transfer appropriate processing architecture where memory encoding and memory retrieval share the same generative operation. At a neural level, our framework consequently predicts that higher similarity between encoding and retrieval will mark episodic memory success [70, 71]. Crucially, generative approximation in memory encoding is not incompatible with the observation of encoding and retrieval modes in the brain [72, 73]. In our model, there is still a fundamental distinction between approximation of current input during encoding and the reinstatement of previous memories during retrieval; the shared generation process does not preclude meaningful differences in other correlates of memory formation and retrieval.

Our research is situated in a growing body of computational modeling work that leverages large generative AI models as models of human memory [26, 74–76]. In line with the complementary learning systems framework [11] our model separates the storage of latent representations in a fast-learning hippocampal system from the neocortical weights that capture statistical knowledge. Importantly, however, we make a novel proposition about how episodic memories are represented across these two complementary systems: our model effectively learns episodic memory representations with complementary components in a way that is distributed across the entire system and it uses a single generation process to find representations at encoding and to reinstate them during retrieval.

Future research could extend this principle in several ways. We have yet to address the implications of a generative approximation framework for consolidation and replay [77]; compare: [26, 78]. Future models may also incorporate sequence generation, which is a domain in which modern AI models excel [79]. Generative encoding of full sequence models could also be extended to naturalistic videos [80], where naturalistic events may emerge as key statistical units from prediction [81, 82]. Our model’s architecture is further limited to a modern Hopfield network, which we use as a stand-in for a hippocampus that stores starting points for regeneration. Future work could merge the generative approximation framework with biologically plausible hippocampal architectures (e.g., [83]).

More broadly, these results suggest that episodic encoding can be understood not as storing a copy of the past, but as searching for a starting point from which the past can be rebuilt. Such an account is parsimonious, because it does not require symmetric back-projections; it is flexible, because multiple generative solutions can support equivalent reinstatement, and it is adaptive, because it frontloads computation to encoding, when time and resources are available and depth can be tuned to expected retrieval demands.

## Methods

### Computational Framework

Our modeling approach rests on the idea that memory encoding and retrieval can be understood as a unified generative process [29] with complementary storage components [11]. We use a pretrained image-generation model whose learned weights capture the statistical regularities of natural images as a computational analogue of cortical knowledge. Encoding an experience corresponds to inverting this generative model, transforming a rich sensory input into a progressively more abstract latent code that is stored in a model of hippocampus. Retrieval corresponds to applying the generative model forward from that code, reinstating the original experience with fidelity that depends on how deeply it was encoded in the first place. We model hippocampus as a separate associative memory module that serves to store the latent resulting from the inversion process. The depth of the inversion process, which is the number of approximation steps, serves as a central variable throughout this work, providing a continuous measure of encoding depth.

Critically, this framework is not tied to any single generative architecture. Different families of generative models learn internal representations of natural image structure, and many admit inversion procedures that recover latent codes from observed images. These inversion methods differ in mechanism: diffusion models use trajectory-based inversion that reverses the forward noising process step by step, whereas generative adversarial networks (GANs) typically use optimization-based inversion that holds network weights fixed and iteratively updates the latent input to minimize reconstruction error [39, 84]. Despite this mechanistic difference, both approaches implement the same computational principle: analysis by synthesis in which a latent representation is found such that generation through the model’s prior knowledge approximates the sensory input [29].

### Generative approximation via fixed feedback in a toy model architecture

We used a multilayer perceptron (MLP), which is a feedforward neural network composed of successive learned linear transformations interleaved with nonlinear activation functions, to map a one-dimensional latent variable (angle *θ*) to an RGB color triplet. The input layer projected *θ* onto its sine and cosine components; these two values fed into an 8-unit hidden layer with rectified linear unit (ReLU) activations; and a 3-unit output layer where sigmoid activations produced RGB values in [0, 1].

The model was trained on 32,000 color samples drawn uniformly from a continuous hue wheel, with hue determined by *θ* and saturation and value both fixed at 1.0 in HSV space. We minimized mean squared error between predicted and target RGB values using stochastic gradient descent (learning rate 0.002, batch size 256, 300 epochs). To verify that the network captured the circular topology of the color space, we applied t-SNE [85] to the 8-dimensional hidden activations for 1,000 uniformly spaced angles, using circular initialization to preserve topology.

To recover the latent angle corresponding to a given target color, we implemented analysis by synthesis through gradient-based optimization: at each step, the current angle estimate *θ* is passed through the frozen generator to produce an RGB output, the mean squared error between this output and the target color is computed, and the gradient of this error with respect to *θ* is obtained via backpropagation through the frozen network weights. Only the latent angle is then updated:

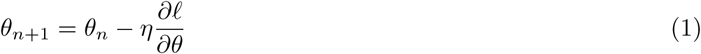

where *η* is the learning rate and *ℓ* = 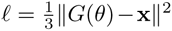 is the mean squared reconstruction error between the generator output *G*(*θ*) and the target color **x**. This procedure is directly analogous to the optimization-based inversion used with StyleGAN-XL (see below) and to analysis by synthesis more broadly [29]: the system iteratively refines a latent representation by evaluating how well generation from the latent matches the sensory input. Because the loss landscape may contain local minima, each encoding attempt was initialized from multiple random starting angles (restarts), and the solution with the lowest final reconstruction error was selected.

### Latent Diffusion Models and Denoising Diffusion Implicit Model (DDIM) Inversion

Latent diffusion models are generative neural networks that learn to synthesize images by iteratively denoising random noise into structured visual content [35, 86]. Training proceeds by gradually corrupting images with Gaussian noise according to a fixed schedule and then teaching a neural network (the “denoiser”) to reverse each corruption step. After training, the model can generate new images by starting from pure noise and applying the learned denoising steps in sequence. In our framework, the pretrained denoiser’s weights embody a rich prior over natural image structure, which we envision as a computational analogue of the cortical statistical knowledge that shapes how new experiences are encoded into memory.

For our main analyses, we used Stable Diffusion v1.5 [35], a widely adopted latent diffusion model trained on a large corpus of natural images. Rather than operating directly on high-dimensional pixel arrays, this model first compresses each image into a compact latent representation using a Variational Autoencoder (VAE, [38]), a neural network that learns to map high-dimensional images into a lower-dimensional latent space and back. The term “latent” refers to the fact that all subsequent operations, both the noising process during training and the inversion and reconstruction procedures during our experiments, are performed in this compressed representational space (for example, a 4 *×* 8 *×* 8 latent tensor for a 3 *×* 64 *×* 64 image; image sizes are given for each experiment below).

To learn a distributed representation during encoding, we applied a Denoising Diffusion Implicit Model (DDIM) inversion [57], which is a computationally effective way to reverse the generative sampling process. Where standard diffusion sampling starts from noise and iteratively denoises to produce an image, DDIM inversion starts from a clean latent and iteratively adds structured noise, producing a trajectory of increasingly abstract representations. Concretely, given a latent state **z***_t_* at timestep *t*, the model first predicts the noise component ***ɛ****_θ_*(**z***_t_, t* + 1), and the next (noisier) state is computed as:

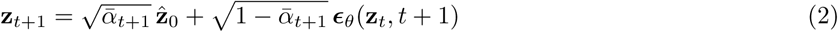

where *ᾱ_t_* is the cumulative product of the noise schedule coefficients [86], and ***ɛ****_θ_* is the learned denoiser. For each encoding depth, a separate inversion was run on a grid of timesteps spaced evenly between 0 and *T*, so that every inversion ended at the noise level *t* = *T* and the encoding depth set the number of steps into which this trajectory was divided. This inversion constitutes the model’s encoding process.

To retrieve (reinstate) a stored memory, we reversed the trajectory: starting from an inverted latent **z***_T_* and applying deterministic DDIM sampling back toward image space. At each step, the model first estimates the fully denoised image **z**^_0_, then interpolates to the next cleaner state:

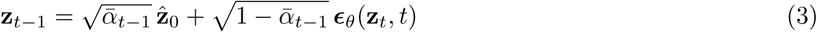

This reconstruction is exact (stochasticity parameter *σ* = 0) when the full trajectory is preserved, and degrades gracefully as inversion depth decreases. All experiments used a uniform linear timestep schedule over the diffusion horizon [0*, T*], with the number of steps *S* varying by experiment (specified below). We disabled Classifier-Free Guidance (CFG [87]) throughout. CFG is a technique used during generation to steer the model’s output toward a specified text description, disabling CFG (guidance scale = 1) is equivalent to unconditional generation, which we preferred because our goal was to study the model’s intrinsic generative process over natural images, not its ability to follow text instructions. For the quantitative analyses, an empty text prompt served as a stand-in for the conditioning input unless specified otherwise.

### StyleGAN-XL image inversion

DDIM inversion is computationally efficient because the number of encoding steps is fixed a priori and each step is a deterministic transformation. To demonstrate that the generative approximation framework does not depend on this specific inversion mechanism, we additionally implemented encoding via StyleGAN-XL [36], a generative adversarial network in which inversion proceeds through iterative optimization: at each step, the model generates an image from the current latent code, computes the reconstruction error relative to the target, and updates the latent via gradient descent. While this method requires more computational resources, we included this proof-of-principle in our simulations because we consider it a more plausible model of neural processes. StyleGAN-XL optimization-based inversion also more directly instantiates analysis by synthesis [29], as the system actively searches for a latent representation by repeatedly evaluating how well its generated output matches the sensory input.

StyleGAN-XL synthesizes images by mapping a latent code through style-modulated convolutional layers that control visual features at different spatial scales. We performed inversion in the extended *W*+ latent space [39], where each of the generator’s style-modulated layers receives its own style vector rather than sharing a single latent code. Early-layer style vectors govern coarse structure, middle layers govern mid-level features, and late layers govern fine details. This per-layer control enables high-fidelity reconstruction of real images.

The latent code was initialized from the generator’s mean latent **ŵ** (estimated by averaging mapped representations of a large random sample) and optimized to reconstruct the target image by minimizing:

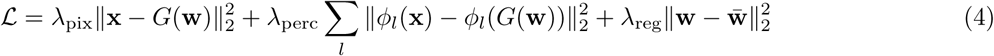

where *ϕ_l_* denotes intermediate feature activations of a pretrained VGG-16 network [88]. The first term enforces pixel-level fidelity, the second captures mid-level perceptual similarity, and the third regularizes against degenerate solutions far from the learned manifold. Weights were set to *λ*_pix_ = 1, *λ*_perc_ = 0.8, *λ*_reg_ = 0.01. We optimized using Adam [89] with learning rate 0.01 for 1,000, 3,000, and 10,000 gradient steps, representing increasing encoding depth. Encoding depth in this context corresponds directly to the number of optimization steps: more steps yield progressively more accurate latent codes and higher-fidelity reconstructions, paralleling the depth-dependent effects observed with DDIM inversion.

### Memory Retrieval

The second component of our model architecture is episodic memory storage of latent representations. When appropriate, we implemented memory storage in our experiments with a Modern Hopfield Network (MHN) operationalizing retrieval as a similarity match in latent space. Given a probe latent representation and a set of stored memory traces, the stored pattern with the highest cosine similarity to the probe was retrieved as a memory. This retrieval metric implements retrieval dynamics of the MHN, which are equivalent to a softmax-weighted attention operation over dot-product similarities between the query and stored patterns, the same similarity computation that underlies cosine similarity for normalized vectors. When stored patterns are sufficiently well-separated in this similarity space, the softmax further converges to the single pattern most similar to the query [90].

### Experiment 1: Generative Approximation on a Color Wheel

To contrast weak and strong encoding, we ran two conditions with the toy model of color. The conditions differed only in the number of optimization steps: weak encoding used 10 gradient steps with 3 random restarts; strong encoding used 200 gradient steps with 3 restarts (both: learning rate *η* = 0.05). To ensure a paired comparison, both conditions were initialized from the same set of random starting angles on each run. We repeated each condition across 25 independent paired runs on a target hue of 25*^◦^* (RGB: [1.0, 0.42, 0.0]). The number of gradient steps hence operationalized encoding depth. We tested the difference in approximation accuracy statistically by computing the absolute angular error between the recovered angle and the true target angle (25*^◦^*) for each run. Because angular errors are bounded and potentially non-normally distributed, we compared weak and strong conditions using a one-tailed paired Wilcoxon signed-rank test (testing the directional hypothesis that strong encoding produces smaller angular errors than weak encoding). We computed a paired test because both conditions shared the same starting angles on each run.

### Experiment 2: Category-Structured Retrieval and Representational Similarity

To characterize the representational format of latent representations and assess their viability for episodic memory tasks we tracked how similarity among stored items evolves across inversion depth and reconstruction steps. To this end we implemented (1) a forced-choice retrieval test quantifying category discriminability and similarity of stored memories, and (2) a representational similarity analysis characterizing the geometry of the latent space.

#### Stimuli

For the in-domain analysis, we selected 300 images from Tiny ImageNet [91], spanning 6 object categories with 50 images each. Images were resized to 64 *×* 64 pixels and normalized to [*−*1, 1] per channel. For out-of-sample evaluation, we assembled 60 images from three categories (bicycles, cats, flowers, 20 images each) collected from Reddit posts dated after December 2023, well beyond the Stable Diffusion v1.5 training-data cutoff date. This ensured that out-of-sample images were unlikely to have appeared during model pretraining.

#### Reconstruction visualization

To visualize how encoding depth affects reconstruction fidelity for single images, we encoded several example images at various depths. For Stable Diffusion, test images were encoded via DDIM inversion at 10 different depths (*S ∈ {*2, 4, 8, 16, 32, 64, 128, 256, 512, 999*}* steps) at a resolution of 512 *×* 512 pixels and reconstructed from each terminal latent by running DDIM sampling with the same number of steps. For StyleGAN-XL, the same images were encoded with 1,000, 3,000 and 10,000 optimization steps and regenerated from the resulting latent code.

#### Category retrieval experiment

The full memory pipeline was implemented with Stable Diffusion as an image generation model and a MHN storage. An image pair from two different image classes was first encoded via image inversion and the resulting latent starting points were stored in the MHN. The latent of a different probe image from one of the categories was then computed and a memory was retrieved by selecting the stored latent with the highest cosine similarity to the probe’s latent representation. Retrieval success was determined based on the category-match between the retrieved image and the probe. This experiment was repeated at 100 encoding depths (10, 20, . . ., 990 and 999 steps).

To compute latent representations in Stable Diffusion, each of the 300 Tiny ImageNet images (6 classes *×* 50 images) was encoded via a separate DDIM inversion at each encoding depth, and the endpoint of each inversion was retained. To compare latent representations in a triplet comparison test, we sampled the anchor image *a* from class *c_a_*, a second image *p* from the same class *c_a_*, and a third image *n* from a different class *c_n_*. The flattened latent vectors were compared via cosine similarity:

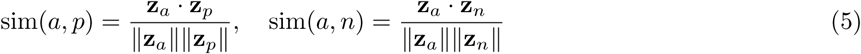

Trials were scored as a same-class retrieval if sim(*a, p*) *>* sim(*a, n*), and as a different-class outcome if the inequality reversed; ties were excluded. We performed 1,000 random triplet comparisons per encoding depth for Tiny ImageNet (500 for the out-of-sample set) in each of five independent runs. For visualization, retrieval-proportion curves were averaged across runs.

#### Representational similarity analysis

To characterize the geometry of latent representations directly, we performed time-resolved representational similarity analysis (RSA [53]). RSA quantifies the relational structure among a set of stimuli by computing all pairwise similarities and asking how that structure changes across encoding and retrieval depth. The analysis was run on five non-overlapping image sets, each consisting of 300 Tiny ImageNet images (6 classes *×* 50 images; 30 classes in total). The first set was the one used in the category-retrieval analysis above. Each set was inverted separately at each encoding depth and reconstructed once from the deepest inversion state.

At each encoding depth and at every tenth step of the reconstruction from the deepest inversion state (steps 1, 11, . . ., 991 and 999 of DDIM sampling), we computed a 300 *×* 300 cosine similarity matrix **S** over all image latents. To exclude trivially perfect self-matches, diagonal entries were omitted from all summary statistics.

Two summary measures were extracted at each timestep. Within-class similarity *s̄*_within_ is the mean of the strictly upper-triangular entries within each of the six 50 *×* 50 diagonal blocks corresponding to the six categories:

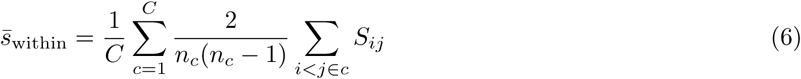

where *C* is the number of classes and *n_c_* is the number of images per class. Between-class similarity *s̄*_between_ is the mean of all entries in the *C*(*C −* 1)*/*2 off-diagonal cross-category blocks:

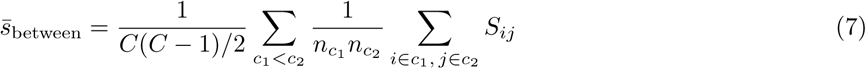

The separation Δ = *s̄*_within_ *− s̄*_between_ indexes the category information carried by the latent representations at each point in the trajectory. These curves were computed across the five image sets and are reported as the mean *±* standard error of the mean (SEM). Separate plots were generated for the inversion phase (Fig. 2G) and the reconstruction phase (Fig. 2I), with Δ on a second axis.

#### Statistics

For the retrieval experiment, the same-category proportion was tested against chance with a one-sided binomial test on the counts pooled over the five runs and all encoding depths. Whether this proportion changed with encoding depth was assessed with the Pearson correlation between depth and same-category proportion, its Bayes factor *BF*_01_ (stretched-beta prior of width 1), and a comparison of the first and last third of encoding depths. For the similarity analysis, within- and between-category similarity were compared at each stage of the trajectory with a one-sided paired *t*-test across the five image sets (Benjamini–Hochberg corrected); monotonic trends during reconstruction were assessed with Spearman correlations between step and the mean similarity, and the change of the within-minus-between difference was estimated by linear regression on step. Whether this difference stayed constant across encoding depths 50 to 999 was tested with two one-sided tests on the linear change of each image set over these depths, against equivalence bounds of *±*10% of the mean difference. Cohen’s *d* denotes the mean of the within-minus-between difference across the five image sets divided by its standard deviation.

#### Experiment 3: Mnemonic Similarity Task Simulation

##### Stimuli and trial construction

We used the standard MST stimulus set, which consists of image pairs organized into five levels of perceptual similarity (level 1 = most similar to the studied item, level 5 = most distinct); the full set comprises 6 MST subsets. Images were resized to 128 *×* 128 pixels, yielding a 4 *×* 16 *×* 16 latent, and pixel values were normalized to [*−*1, 1].

Stimuli were organized into batches of 128 image pairs. Within each batch, the first images of 64 pairs served as study items. At test, 192 probes were constructed from three sources: 64 old probes (exact repetitions of study items), 64 lure probes (the paired counterparts of the study items, spanning five similarity levels), and 64 new probes (the first images of the remaining 64 pairs, which had no studied counterpart). The test phase of each batch therefore matched the behavioral MST (64 repeats, 64 lures, 64 foils); every studied item was probed both once as a repeat and once as a lure. Probes carried a 7-level label: 0 = old, 1–5 = lure levels 1 through 5, and 6 = new. The 1,152 pairings of the six MST sets were repeated three times, reshuffling on each pass which pairs were studied and which served as new probes, resulting in 9 batches per pass (27 total), with 1,728 study presentations and 5,184 test probes per encoding-depth condition.

##### Encoding and retrieval

To simulate different levels of encoding depth, we encoded the study items at three DDIM inversion depths: 999 steps (strong encoding), 20 steps (moderate encoding), and 5 steps (weak encoding). Each study item’s final inversion latent (the latent state reached at the end of the image inversion trajectory) served as its stored memory trace. To simulate retrieval, we reconstructed each memory by running DDIM sampling from the stored latent using the same number of steps as the original encoding. The reconstructed latent was then flattened into a 1024-dimensional vector. To determine retrieval accuracy, we quantified the cosine similarity between the reconstructed memory vector (a remembered image) and the test probe’s initial VAE-encoded latent representation (sensory input prior to any diffusion process). Specifically, for each test probe **p** and its corresponding reconstructed memory **m**^, we computed:

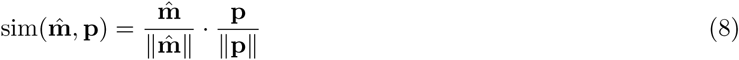

where ∥ · ∥ denotes the *ℓ*_2_ norm. Higher similarity indicates that the reconstructed memory more closely matches the test probe. Repeated and lure probes were compared with the reconstructed memory of their own studied item; new probes, which have no studied counterpart, were compared with the reconstructed memory of another studied item of the same batch.

##### Reconstruction fidelity

To verify that encoding depth controls how well an image is recovered, reconstructions were compared with the studied images in pixel space. The first 256 study images of the stimulus set were fitted into a square with white padding and downsampled to 64 *×* 64 pixels. Each image was inverted and reconstructed at 5, 20 and 999 steps, the reconstructed latent was decoded with the VAE decoder, and the structural similarity index (SSIM [43]) between the decoded and the original image was computed. Deep (999 steps) and shallow (5 steps) encoding were compared with a paired Wilcoxon signed-rank test across images. SSIM served as a measure of image fidelity only; mnemonic discrimination was assessed on the cosine similarity between latent representations (Eq. 8), because retrieval in the model operates on latents.

##### Normalization and similarity comparison

To compare memory performance across the three encoding-depth conditions, raw similarity scores were normalized to a range between 0 and 1 for each condition. Specifically, for each condition, the mean cosine similarity of old probes (*µ*_old_) and the mean of new probes (*µ*_new_) served as anchors. Each stimulus category’s mean similarity *s̄_k_* was linearly transformed:

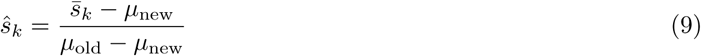

This produces a scale of recognition strength based on similarity to old items, which is consistent with the convention used in the MST literature, where the probability of endorsing an item as old *p*(old|item) is measured.

##### Visualization of differences

Finally, for visualization purposes only, we expressed the lure similarity gradient relative to a linear increase in the deepest encoding depth condition (999 steps), i.e., the increase in similarity in the deepest condition (999 steps) was used as a reference that was set to increase linearly. Specifically, equally spaced target values *{v_k_}* were defined for the 7 stimulus classes. For each lure level *k*, a scaling factor was computed from the 999-step condition:

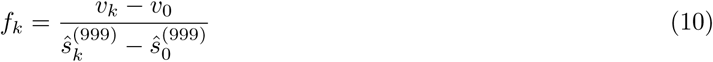

These five factors were then applied to the corresponding lure levels in the 20-step and 5-step conditions effectively multiplying each lure level in those conditions with a constant derived from the deepest conditions:

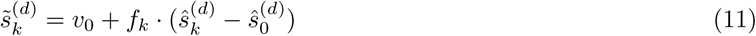

This calibration preserves relative deviations from the deep-encoding reference while keeping endpoints aligned, facilitating direct visual comparison of how encoding depth reshapes the lure discrimination gradient.

##### Statistics

Because each stimulus pair could appear in more than one pass, we first identified repeated presentations of the same pair from their initial VAE latent and averaged their similarity values, so that every pair contributed one value per depth condition (about 200 pairs per lure level and depth). To test the ordering of stimulus classes, we compared raw cosine similarities of old probes against all lure probes, and of all lure probes against new probes, within each depth condition using Welch’s *t*-test. To test the effect of encoding depth we used the anchored values *ŝ* (Eq. 9). Because the same pairs were tested at all three depths, we treated depth as a within-pair factor: at each lure level we ran a repeated-measures ANOVA over depth on the pairs that were lures at all three depths, with Greenhouse–Geisser correction, and paired *t*-tests for each pairwise contrast on the pairs shared by the two depths.

#### Rational Analysis of Encoding-Depth

In our framework, deeper encoding improves discrimination but takes longer. We formalized this tradeoff over 18 encoding depths between 3 and 80 DDIM steps (3 to 10 in steps of one, 12, 15, 20, 25, 30, 35, 40, 50, 60 and 80) by defining a combined reward function that balances discrimination against the computational cost of deep encoding. For each depth *d* and lure bin *k* we considered the lure items’ normalized similarity score 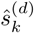, which are values between 0 (lure no more similar to the memory than a novel item) and 1 (lure as similar to the memory as an exact repetition). We then defined discrimination reward for lures as the complement:

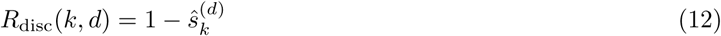

Higher values in discrimination reward denote that the model’s retrieved trace is more distinct from the lure and the memory is more useful. This reward rose with depth because deeper encoding produced cleaner traces and larger separation from confusable lures. Because deeper encoding requires more computation, we then defined an efficiency reward that decreases linearly with depth:

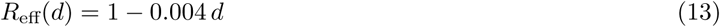

The efficiency schedule assigns a reward of 0.99 to the shallowest depth (3 steps) and of 0.68 to the deepest (80 steps), effectively implementing a reward for less computation. We computed the outcome of this schedule for all five lure bins across encoding depths. Finally, we multiplied discrimination and efficiency reward to obtain a total reward:

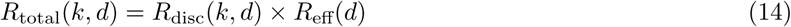

We then identified, for each lure bin, the sampled depth with the highest total reward, demonstrating how the trade-off between representation and computation can shift optimal encoding policies if computation becomes associated with increasing cost. Finally, we computed the Spearman rank correlation between lure indices and each lure’s depth of maximum total reward.

#### Information Distribution Across the Network

##### Mutual information estimation

To quantify how information about the original image is distributed between the latent representation and the generator weights at each encoding depth, we estimated mutual information (MI) between pairs of latent representations using InfoNCE [92]. InfoNCE recasts the estimation of MI as a classification problem that yields a lower bound on MI in high-dimensional latent spaces, where direct MI estimation is intractable.

Specifically, given a batch of *N* matched latent pairs *{*(**z***_i_,* **z***^′^_i_*)*}* drawn from the joint distribution, we *ℓ*_2_-normalized each vector and computed a cosine similarity matrix **C**. Each row of this matrix contains one “correct” pairing (the diagonal element *C_ii_*) and *N −* 1 incorrect pairings. We then applied a standard cross-entropy loss treating the diagonal as the target class:

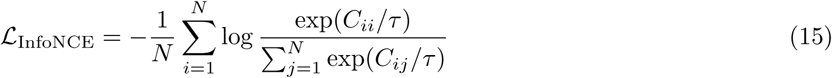

We used *τ* = 1, that is, we computed cross-entropy loss without temperature scaling. Because cosine similarities are bounded between *−*1 and 1, the maximum estimate under *τ* = 1 falls below the log *N* ceiling. Absolute values are therefore conservative, and we interpret how the estimate changes across encoding depth, rather than its magnitude.

If **z** and **z***^′^* share no information, the classifier performs at chance and the loss equals log *N* . If they share maximal information, the loss approaches zero. The MI lower bound is the gap between these extremes:

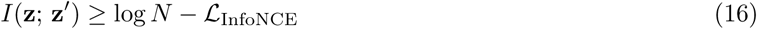

Each estimate used a batch of *N* = 80 images, capping its upper bound of MI at log 80 *≈* 4.38 nats. To compute MI across images, each image was first processed through *S* steps of DDIM inversion followed by *S* steps of reconstruction. We extracted three anchor latents from the trajectory: The “start” latent at the first VAE step, representing the original image encoding, the “deepest” latent at inversion step *S*, representing the point of final approximation; and the final “reconstructed” latent, representing the retrieved image. From these anchors we tracked two MI values across encoding depths. The MI estimate between the start and deepest latent tracks how much information about the original image remains detectable by the similarity-based readout as encoding progresses, which is expected to decline with deeper encoding. The MI estimate between start and reconstructed latent tracks how much of that information becomes detectable again after the full encode-then-retrieve cycle. The crossover point between decaying information with the latent and increasing information with the reconstructed image marks the encoding depth at which more information about the image is recovered from the weights of the network than is detectable in latent space itself. While the absolute MI is not always comparable between different image modalities because images within a sub-category are more homogeneous and distinguishing them can be harder, we leverage the crossover point as an indicator of how quickly information can be approximated through the network. It denotes the number of encoding steps needed until reconstruction quality exceeds information accessible from the latent representation alone. We sampled encoding depths on a grid of 29 points between 3 and 305 steps (3, 4, 5, 6, 8, 10, 12, 15, 20, 25, 30, 35, 40, 45, 50, 60 and 65 steps, followed by 20-step spacing from 85 to 305 steps). To assess reliability, every condition was run in five independent iterations, each drawing a new batch of 80 images.

To detect the crossover, we computed the difference Δ = MI_recon_ *−* MI_latent_ at each sampled depth and placed the crossover by linear interpolation between the two adjacent depths at which Δ changed sign. In each of the 45 instances (9 conditions with 5 iterations), curves crossed exactly once within the sampled range (individual crossovers from 15.2 to 155.0 steps). Reported crossover depths are means of the five iterations in each condition.

#### Experiment 4: Domain Expertise

##### Domain-specific finetuning

To model domain-specific expertise, we modified the base Stable Diffusion v1.5 model with Low-Rank Adaptation (LoRA [51]). LoRA freezes the original model weights and injects pairs of small trainable matrices into each attention layer, so that the adapted weight matrix becomes *W^′^* = *W* + *BA* where *B ∈* R*^d×r^* and *A ∈* R*^r×d^* with rank *r ≪ d*. This modifies only a fraction of the total parameters while shifting the model’s generative distribution toward a target domain. We trained two domain-tuned variants with identical hyperparameters, differing only in their training image source and in a single static caption token. The cartoon variant was trained on 500 images drawn from the Danbooru anime archive and used the static caption “sks cartoon”; the pixel-art variant was trained on 500 images drawn from the public DiffusionDB pixel-art subset and used the static caption “sks pixel”.

The rare “sks ” prefix avoids collision with the semantic concepts the base model already associates with the words “cartoon” and “pixel”. Training images were resized to 128 *×* 128 to match the evaluation resolution, applying resize and tensor conversion only, with no per-channel normalization. LoRA adapters were configured at rank 32 with alpha 32 and injected into the UNet attention projections (to q, to k, to v, to out.0), the transformer block projections (proj in, proj out), the ResNet block convolutions (conv1, conv2, conv shortcut), and the CLIP text-encoder attention projections (q proj, k proj, v proj, out proj). Each LoRA was trained for 4,000 optimizer steps at a per-device batch size of 4 with gradient accumulation over 2 mini-batches, using a learning rate of 1 *×* 10*^−^*^4^ for the UNet adapters and 5 *×* 10*^−^*^5^ for the text-encoder adapters under a cosine schedule, with the denoising loss reweighted by the Min-SNR scheme [93] (snr gamma = 5.0). Each set of LoRA weights was then fused into the base model’s U-Net and text encoder prior to latent extraction.

##### Experimental design

We evaluated all nine pairwise combinations of three models (the untuned base, the cartoon-tuned variant, and the pixel-tuned variant) crossed with three image domains: natural photographs drawn from Tiny ImageNet [91], cartoon images drawn from the Danbooru archive, and pixel-art images drawn from the DiffusionDB pixel-art subset. Two of the nine conditions represent matched expertise, the cartoon model processing cartoon images and the pixel model processing pixel art; the remaining seven conditions served as mismatched or untuned controls. Within each condition we ran five independent replicates, each drawing a fresh batch of 80 images from the domain pool, yielding 45 mutual-information curves in total. The prompt presented to the model was bound to the image domain rather than to the LoRA: cartoon images were paired with “sks cartoon”, pixel-art images with “sks pixel”, and natural images with the empty prompt, holding the prompt constant across the three models within each image domain so that any cross-model difference in crossover depth reflects the LoRA-adapted weights rather than a change in prompt.

##### Statistical comparison

To test for MI increases and decreases with increasing encoding depths, we computed a Spearman correlation coefficient between encoding depth and the ranked MI across all encoding depths. To test whether domain-matched models reach the MI crossover earlier than mismatched models, we computed the interaction contrast *T* = (CC + PP) *−* (CP + PC), where CC and PP denote the two matched model-domain pairings and CP and PC the two mismatched pairings, with a negative value indicating earlier crossover for matched pairings. We constructed an empirical null distribution by shuffling the two LoRA model labels within each image-domain pool, independently for the cartoon-image and pixel-art pools, and recomputing *T* for each of 10,000 such reassignments. Shuffling within image domain preserves the substantial baseline differences in crossover depth across domains, which are present even for the untuned base model and would otherwise inflate the null variance; the *p*-value is the fraction of shuffled contrasts at least as extreme as the observed one.

#### Neural Representational Alignment

##### Natural Scenes Dataset

We used the Natural Scenes Dataset (NSD [52]), a large-scale 7-Tesla fMRI resource in which eight participants each viewed up to 10,000 distinct color natural scenes drawn from the COCO image set. Images were presented for 3 seconds followed by a 1-second inter-trial gap, and each image could be repeated up to three times across 30–40 scanning sessions. We analyzed Sessions 1–3 for all eight participants, yielding approximately 2,250 stimulus–response pairs per participant and 11,593 unique images across the cohort. We computed matched latent representations for the NSD images, such that every model latent vector could be paired with the fMRI pattern evoked by the same image in the same participant.

##### Regions of interest

We defined 16 regions of interest (ROIs) spanning early visual cortex through hippocampal subfields. These ROIs were organized into four functional groups. The Early Visual group comprised the ventral and dorsal subdivisions of V1 (V1v, V1d). The Mid-level Visual group comprised the ventral and dorsal subdivisions of V2 (V2v, V2d). The Higher Visual group comprised the ventral and dorsal subdivisions of V3 together with area hV4 (V3v, V3d, hV4). The MTL group comprised parahippocampal cortex (PHC), perirhinal cortex areas 35 and 36 (PRC35, PRC36), entorhinal cortex (ERC), hippocampal subfields CA1, CA2, and CA3, dentate gyrus (DG), and the hippocampal tail (HT). ROI masks were obtained from the NSD subject-specific volumetric atlases in native functional space (1-mm resolution). For each participant, session, and ROI, we then extracted single-trial beta estimates from the GLMdenoise-derived response volumes [94], and *z*-scored voxel responses within each ROI, so that subsequent similarity analyses reflected multi-voxel activation patterns rather than overall signal magnitude.

##### Model representations

Each NSD stimulus image was resized to 64 *×* 64 pixels using bilinear interpolation with antialiasing. This resolution was chosen because Stable Diffusion’s VAE encoder produces 8*×* spatial downsampling, so a 64 *×* 64 input yields a 4 *×* 8 *×* 8-dimensional latent code per timestep, compact enough to process the full multi-subject NSD stimulus set through 999-step DDIM inversion and reconstruction on a single GPU, while preserving the spatial layout that carries scene-level information.

Each resized image was encoded via the VAE and passed through 999-step DDIM inversion to obtain a noise-level endpoint, then reconstructed via 999-step DDIM sampling.

We retained 34 latent states spaced 30 steps apart from the reconstruction trajectory for the reinstatement analysis. The latent starting point was represented by the endpoint of a separate 999-step inversion.

Each sampled latent had a shape of 4 *×* 8 *×* 8 (channels *×* height *×* width) and was flattened into a 256-dimensional feature vector. This produced, for each sampled depth *d*, a trial-by-feature matrix **X***_d_ ∈* R*^n×^*^256^ where *n* is the number of stimulus presentations for that participant.

##### Centered Kernel Alignment

We quantified model–brain correspondence using linear Centered Kernel Alignment [54]. CKA compares the pattern of pairwise relationships among stimuli (compare: [53]). If two images that evoke similar voxel patterns in V1 also produce similar diffusion latents at some depth, CKA will be high for that depth–ROI pair. We used CKA, because it is invariant to rotation and isotropic scaling of either representation, making it suitable for comparing spaces with different dimensionalities and arbitrary coordinate systems.

Formally, for a given depth *d* and ROI *r*, let **X** and **Y** denote the trial-matched model and neural feature matrices (*p* model features; *q* number of voxels in that ROI). The computation proceeded in four steps:

### Column-centering

Each feature column of **X** and **Y** was mean-subtracted, removing the mean activation across trials so that only stimulus-driven variance remained.

### Gram matrices

We computed trial-by-trial similarity matrices **K***_X_* = **XX***^⊤^* and **K***_Y_* = **YY***^⊤^*. Each entry *K_X,ij_* captures the dot-product similarity between the model’s representations of trials *i* and *j*; each entry *K_Y,ij_* captures the corresponding neural similarity.

### Double-centerin

Both Gram matrices were then centered using **K̂** = **HKH** where 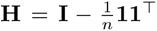11^T^. This removes row and column means, ensuring that the subsequent similarity measure is not inflated by shared global structure.

### HSIC and normalization

The Hilbert–Schmidt Independence Criterion (HSIC) quantifies the agreement between the two centered similarity structures:

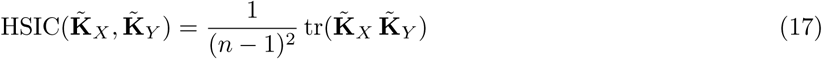

Linear CKA normalizes HSIC by the geometric mean of each space’s self-similarity, yielding a score between 0 and 1:

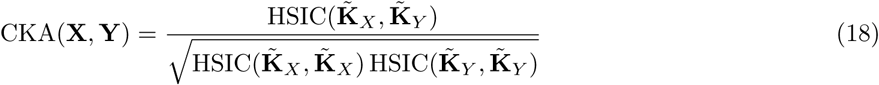

A CKA of 1 means the two spaces impose identical similarity rankings over stimuli; a CKA of 0 means the rankings are unrelated.

This procedure yielded one CKA score for every combination of participant (*n* = 8), ROI (16 regions), and model state (the starting point and 34 reconstruction states), producing a state *×* ROI heatmap per participant.

#### Significance of the model–brain correspondence

Before relating these correspondences to the cortical hierarchy, we asked whether the CKA values reflected image-by-image correspondence rather than incidental similarity between the two representational spaces. For each depth–ROI combination we built a chance-level distribution by shuffling, independently within each participant, which trial’s fMRI response was paired with which model latent and recomputing CKA; this left each representation’s pattern of pairwise similarities intact but broke the link between individual images and neural responses. For each of 1,000 such shuffles we averaged the resulting CKA across the eight participants, resulting in a null distribution of eight-participant-averages. We then took the one-sided permutation *p*-value as the fraction of these shuffled averages as large as or larger than the observed average, with the conventional add-one correction so that its smallest attainable value was 1*/*1001 *≈* 0.001. We used a one-sided test because only above-chance correspondence is meaningful. Because this produced one test per depth–ROI combination (35 model states *×* 16 ROIs = 560 tests), we controlled the false discovery rate across all 560 combinations with the Benjamini–Hochberg procedure at *q* = 0.05, treating a combination as significant only when its *p*-value fell below the adjusted threshold rather than rescaling the *p*-values themselves.

### Statistical comparison of hierarchies

All statistical tests were two-tailed with *α* = 0.05, except for tests of directional hypotheses, which were one-sided. We report exact *p*-values except for permutation tests, whose resolution is bounded by the number of shuffles and which are therefore reported as bounds (e.g., *p <* 0.001). Analyses were conducted in Python 3.11 using SciPy 1.15 [95] and Pingouin 0.7. To test the relationship between the cortical hierarchy and reconstruction step in the model, we assigned hierarchy ranks to the 16 ROIs according to the cortical processing scheme of Felleman and Van Essen [56], with sub-regions occupying the same hierarchical level sharing a single rank rather than being arbitrarily ordered within that level: V1v and V1d were both assigned rank 1; V2v and V2d rank 2; V3v, V3d, and hV4 rank 3; PHC rank 4; PRC35 and PRC36 rank 5; ERC rank 6; and CA1, CA2, CA3, DG, and HT rank 7. We then correlated each ROI’s mean peak-CKA depth with its hierarchy rank via Spearman rank correlation, across the steps of reconstruction from the deepest latent at 999 inversion depth. Peak depth was defined, for each participant and ROI, as the sampled depth with the highest CKA. For the correlation with hierarchy rank, peak depths were averaged across participants, and the significance of the correlation was assessed by permuting the rank assignments 10,000 times. For the comparison between V1 and MTL, peak depths were averaged across the ROIs of each group within each participant and compared across participants with a paired *t*-test and a Wilcoxon signed-rank test. The peak markers in Fig. 5 show the peak of the group-average CKA curve of each ROI, which differs from the mean of the participants’ peaks by less than one sampling interval (30 steps); the colors rank the ROIs by this peak, with ties broken by the mean of the participants’ peaks.

### Large language model and AI use

The authors used a large language model (Claude Fable 5.1, Anthropic) to audit and review analysis code and for copyediting of the manuscript text. The model did not generate analyses, results, or scientific content. The authors take full responsibility for the work.

## Data availability

No new data were collected for this study. The Natural Scenes Dataset [52] is available at http://naturalscenesdataset.org. The stimulus sets of the Mnemonic Similarity Task [41] are available at https://github.com/celstark/MST. Tiny ImageNet [91] is available at http://cs231n.stanford.edu/tiny-imagenet-200.zip. Cartoon images were taken from the Tagged Anime Illustrations dataset, a subset of Danbooru2017 (https://www.kaggle.com/datasets/mylesoneill/tagged-anime-illustrations), and pixel-art images from the pixel-art subset of DiffusionDB (https://huggingface.co/datasets/jainr3/diffusiondb-pixelart). The weights of Stable Diffusion v1.5 [35] are available at https://huggingface.co/stable-diffusion-v1-5/stable-diffusion-v1-5 and those of StyleGAN-XL [36] at https://github.com/autonomousvision/stylegan-xl. The images collected from Reddit are available from the corresponding author upon request. The data underlying all statistics and figures are provided with the code.

## Code availability

The code for all simulations and analyses is available at Zenodo (https://doi.org/10.5281/zenodo.23093152).

## Acknowledgements

We thank T. Gureckis, A. Goldstein, C. Kerŕen, and M. Schottdorf for comments on the manuscript, and C. Iyer for helpful suggestions.

## Author contributions

Conceptualization: H.L., M.M. and S.M., Methodology: H.L., M.M. and S.M., Code: H.L. and S.M., Formal analysis: H.L., Investigation: H.L., Visualization: H.L. and S.M., Writing (original draft): H.L. and S.M., Writing (review & editing): H.L., M.M. and S.M., Supervision: M.M. and S.M.

## References

[1] Nagy, D. G., Orbán, G. & Wu, C. M. Adaptive compression as a unifying framework for episodic and semantic memory. Nature Reviews Psychology 4, 484–498 (2025).

[2] Ahmad, F. N., Moscovitch, M. & Hockley, W. E. Effects of varying presentation time on long-term recognition memory for scenes: Verbatim and gist representations. Memory & Cognition 45, 390–403 (2017).

[3] Craik, F. I. M. & Lockhart, R. S. Levels of processing: A framework for memory research. Journal of Verbal Learning and Verbal Behavior 11, 671–684 (1972).

[4] Greene, N. R. & Naveh-Benjamin, M. The formation of specific and gist associative episodic memory representations during encoding: Effects of rate of presentation. *Journal of Experimental Psychology: Learning*, Memory, and Cognition 49, 1099–1118 (2023).

[5] Korkki, S. M., Richter, F. R. & Simons, J. S. Hippocampal-cortical encoding activity predicts the precision of episodic memory. Journal of Cognitive Neuroscience 33, 2328–2341 (2021).

[6] Scoville, W. B. & Milner, B. Loss of recent memory after bilateral hippocampal lesions. *Journal of Neurology*, Neurosurgery & Psychiatry 20, 11–21 (1957).

[7] Nieh, E. H. et al. Geometry of abstract learned knowledge in the hippocampus. Nature 595, 80–84 (2021).

[8] Ison, M. J., Quian Quiroga, R. & Fried, I. Rapid encoding of new memories by individual neurons in the human brain. Neuron 87, 220–230 (2015).

[9] Quian Quiroga, R., Reddy, L., Kreiman, G., Koch, C. & Fried, I. Invariant visual representation by single neurons in the human brain. Nature 435, 1102–1107 (2005).

[10] Quian Quiroga, R. Concept cells: The building blocks of declarative memory functions. Nature Reviews Neuroscience 13, 587–597 (2012).

[11] McClelland, J. L., McNaughton, B. L. & O’Reilly, R. C. Why there are complementary learning systems in the hippocampus and neocortex: Insights from the successes and failures of connectionist models of learning and memory. Psychological Review 102, 419–457 (1995).

[12] O’Reilly, R. C. & Norman, K. A. Hippocampal and neocortical contributions to memory: Advances in the complementary learning systems framework. Trends in Cognitive Sciences 6, 505–510 (2002).

[13] Norman, K. A. & O’Reilly, R. C. Modeling hippocampal and neocortical contributions to recognition memory: A complementary-learning-systems approach. Psychological Review 110, 611–646 (2003).

[14] Teyler, T. J. & DiScenna, P. The hippocampal memory indexing theory. Behavioral Neuroscience 100, 147–154 (1986).

[15] Danker, J. F. & Anderson, J. R. The ghosts of brain states past: Remembering reactivates the brain regions engaged during encoding. Psychological Bulletin 136, 87–102 (2010).

[16] Johnson, J. D. & Rugg, M. D. Recollection and the reinstatement of encoding-related cortical activity. Cerebral Cortex 17, 2507–2515 (2007).

[17] Kuhl, B. A. & Chun, M. M. Successful remembering elicits event-specific activity patterns in lateral parietal cortex. The Journal of Neuroscience 34, 8051–8060 (2014).

[18] Polyn, S. M., Natu, V. S., Cohen, J. D. & Norman, K. A. Category-specific cortical activity precedes retrieval during memory search. Science 310, 1963–1966 (2005).

[19] Rissman, J. & Wagner, A. D. Distributed representations in memory: Insights from functional brain imaging. Annual Review of Psychology 63, 101–128 (2012).

[20] Rugg, M. D. & Vilberg, K. L. Brain networks underlying episodic memory retrieval. Current Opinion in Neurobiology 23, 255–260 (2013).

[21] Wheeler, M. E., Petersen, S. E. & Buckner, R. L. Memory’s echo: Vivid remembering reactivates sensory-specific cortex. Proceedings of the National Academy of Sciences 97, 11125–11129 (2000).

[22] Rolls, E. T. Hippocampo-cortical and cortico-cortical backprojections. Hippocampus 10, 380–388 (2000).

[23] Rolls, E. T. Pattern separation, completion, and categorisation in the hippocampus and neocortex. Neurobiology of Learning and Memory 129, 4–28 (2016).

[24] Lavenex, P. & Amaral, D. G. Hippocampal-neocortical interaction: A hierarchy of associativity. Hippocampus 10, 420–430 (2000).

[25] Witter, M. P., Wouterlood, F. G., Naber, P. A. & Van Haeften, T. Anatomical organization of the parahippocampal-hippocampal network. Annals of the New York Academy of Sciences 911, 1–24 (2000).

[26] Spens, E. & Burgess, N. A generative model of memory construction and consolidation. Nature Human Behaviour 8, 526–543 (2024).

[27] Nagy, D. G., Torok, B. & Orban, G. Optimal forgetting: Semantic compression of episodic memories. PLOS Computational Biology 16, e1008367 (2020).

[28] O’Reilly, R. C., Bhattacharyya, R., Howard, M. D. & Ketz, N. Complementary learning systems. Cognitive Science 38, 1229–1248 (2014).

[29] Yuille, A. & Kersten, D. Vision as Bayesian inference: Analysis by synthesis? Trends in Cognitive Sciences 10, 301–308 (2006).

[30] Gilboa, A. & Moscovitch, M. No consolidation without representation: Correspondence between neural and psychological representations in recent and remote memory. Neuron 109, 2239–2255 (2021).

[31] Slamecka, N. J. & Graf, P. The generation effect: Delineation of a phenomenon. Journal of Experimental Psychology: Human Learning and Memory 4, 592–604 (1978).

[32] Roediger, H. L. & Karpicke, J. D. Test-enhanced learning: Taking memory tests improves long-term retention. Psychological Science 17, 249–255 (2006).

[33] Gershman, S. J., Fiete, I. & Irie, K. Key-value memory in the brain. Neuron 113, 1694–1707.e1 (2025).

[34] Rorschach, H. *Psychodiagnostik: Methodik und Ergebnisse eines wahrnehmungsdiagnostischen Experiments* (Ernst Bircher, Bern, 1921).

[35] Rombach, R., Blattmann, A., Lorenz, D., Esser, P. & Ommer, B. High-resolution image synthesis with latent diffusion models. Proceedings of the IEEE/CVF Conference on Computer Vision and Pattern Recognition (CVPR), 10684–10695 (2022).

[36] Sauer, A., Schwarz, K. & Geiger, A. StyleGAN-XL: Scaling StyleGAN to large diverse datasets. ACM SIGGRAPH 2022 Conference Proceedings, 1–10 (2022).

[37] Creswell, A. & Bharath, A. A. Inverting the generator of a generative adversarial network. IEEE Transactions on Neural Networks and Learning Systems 30, 1967–1974 (2019).

[38] Kingma, D. P. & Welling, M. *Auto-encoding variational Bayes*. *International Conference on Learning Representations* (2014).

[39] Abdal, R., Qin, Y. & Wonka, P. Image2StyleGAN: How to embed images into the StyleGAN latent space? Proceedings of the IEEE/CVF International Conference on Computer Vision (ICCV), 4431–4440 (2019).

[40] Franch, M. et al. A population code for semantics in human hippocampus. Nature Neuroscience 1–13 (2026).

[41] Stark, S. M., Kirwan, C. B. & Stark, C. E. L. Mnemonic similarity task: A tool for assessing hippocampal integrity. Trends in Cognitive Sciences 23, 938–951 (2019).

[42] Stark, S. M., Yassa, M. A., Lacy, J. W. & Stark, C. E. L. A task to assess behavioral pattern separation (BPS) in humans: Data from healthy aging and mild cognitive impairment. Neuropsychologia 51, 2442–2449 (2013).

[43] Wang, Z., Bovik, A. C., Sheikh, H. R. & Simoncelli, E. P. Image quality assessment: from error visibility to structural similarity. IEEE Transactions on Image Processing 13, 600–612 (2004).

[44] Kellen, D., Winiger, S., Dunn, J. C. & Singmann, H. Testing the foundations of signal detection theory in recognition memory. Psychological Review 128, 1022–1050 (2021).

[45] Beaudry, N. J. & Renner, R. An intuitive proof of the data processing inequality. Quantum Information & Computation 12, 432–441 (2012).

[46] Xu, Y., Zhao, S., Song, J., Stewart, R. & Ermon, S. *A theory of usable information under computational constraints*. *International Conference on Learning Representations* (2020).

[47] Chase, W. G. & Simon, H. A. Perception in chess. Cognitive Psychology **4**, 55–81 (1973).

[48] Gobet, F. & Simon, H. A. Templates in chess memory: A mechanism for recalling several boards. Cognitive Psychology 31, 1–40 (1996).

[49] Wing, E. A., Burles, F., Ryan, J. D. & Gilboa, A. The structure of prior knowledge enhances memory in experts by reducing interference. Proceedings of the National Academy of Sciences 119, e2204172119 (2022).

[50] Wing, E. A., Gilboa, A. & Ryan, J. D. Interactive influences of prior knowledge on episodic memory. iScience 27, 111142 (2024).

[51] Hu, E. J., et al. *LoRA: Low-rank adaptation of large language models*. *International Conference on Learning Representations* (2022).

[52] Allen, E. J. et al. A massive 7T fMRI dataset to bridge cognitive neuroscience and artificial intelligence. Nature Neuroscience 25, 116–126 (2022).

[53] Kriegeskorte, N., Mur, M. & Bandettini, P. A. Representational similarity analysis – connecting the branches of systems neuroscience. Frontiers in Systems Neuroscience 2, 4 (2008).

[54] Kornblith, S., Norouzi, M., Lee, H. & Hinton, G. Similarity of neural network representations revisited. Proceedings of the 36th International Conference on Machine Learning, PMLR 97, 3519–3529 (2019).

[55] Rolls, E. T., Deco, G., Zhang, Y. & Feng, J. Hierarchical organization of the human ventral visual streams revealed with magnetoencephalography. Cerebral Cortex 33, 10686–10701 (2023).

[56] Felleman, D. J. & Van Essen, D. C. Distributed hierarchical processing in the primate cerebral cortex. Cerebral Cortex 1, 1–47 (1991).

[57] Song, J., Meng, C. & Ermon, S. *Denoising diffusion implicit models*. *International Conference on Learning Representations* (2021).

[58] Molitor, R. J., Ko, P. C., Hussey, E. P. & Ally, B. A. Memory-related eye movements challenge behavioral measures of pattern completion and pattern separation. Hippocampus 24, 666–672 (2014).

[59] Zhou, D. et al. A compressed code for memory discrimination. Preprint at 10.1101/2025.10.12.681901 (2025).

[60] Griffiths, T. L., Lieder, F. & Goodman, N. D. Rational use of cognitive resources: Levels of analysis between the computational and the algorithmic. Topics in Cognitive Science 7, 217–229 (2015).

[61] Mattar, M. G. & Daw, N. D. Prioritized memory access explains planning and hippocampal replay. Nature Neuroscience 21, 1609–1617 (2018).

[62] Tse, D. et al. Schema-dependent gene activation and memory encoding in neocortex. Science 333, 891–895 (2011).

[63] Wallace, B., Gokul, A., Ermon, S. & Naik, N. End-to-end diffusion latent optimization improves classifier guidance. Proceedings of the IEEE/CVF International Conference on Computer Vision (ICCV), 7280–7290 (2023).

[64] Ozcelik, F. & VanRullen, R. Natural scene reconstruction from fMRI signals using generative latent diffusion. Scientific Reports 13, 15666 (2023).

[65] Takagi, Y. & Nishimoto, S. High-resolution image reconstruction with latent diffusion models from human brain activity. Proceedings of the IEEE/CVF Conference on Computer Vision and Pattern Recognition (CVPR), 14453–14463 (2023).

[66] Bonnen, T., Yamins, D. L. K. & Wagner, A. D. When the ventral visual stream is not enough: A deep learning account of medial temporal lobe involvement in perception. Neuron 109, 2755–2766.e6 (2021).

[67] Kali, S. & Dayan, P. Off-line replay maintains declarative memories in a model of hippocampal-neocortical interactions. Nature Neuroscience 7, 286–294 (2004).

[68] Huang, S. et al. Hippocampal functions modulate transfer-appropriate cortical representations supporting subsequent memory. The Journal of Neuroscience 44, e1135232023 (2024).

[69] Morris, C. D., Bransford, J. D. & Franks, J. J. Levels of processing versus transfer appropriate processing. Journal of Verbal Learning and Verbal Behavior 16, 519–533 (1977).

[70] Kragel, J. E. et al. Similar patterns of neural activity predict memory function during encoding and retrieval. NeuroImage 155, 60–71 (2017).

[71] Ritchey, M., Wing, E. A., LaBar, K. S. & Cabeza, R. Neural similarity between encoding and retrieval is related to memory via hippocampal interactions. Cerebral Cortex 23, 2818–2828 (2013).

[72] Hasselmo, M. E., Bodeĺon, C. & Wyble, B. P. A proposed function for hippocampal theta rhythm: Separate phases of encoding and retrieval enhance reversal of prior learning. Neural Computation 14, 793–817 (2002).

[73] Smith, D. E., Moore, I. L. & Long, N. M. Temporal context modulates encoding and retrieval of overlapping events. The Journal of Neuroscience 42, 3000–3010 (2022).

[74] D’Alessandro, M., D’Amato, L., Elkano, M., Uriz, M. & Pezzulo, G. GENESIS: A generative model of episodic–semantic interaction. Neuroscience & Biobehavioral Reviews 185, 106627 (2026).

[75] Dong, C. V., Lu, Q., Norman, K. A. & Michelmann, S. Towards large language models with human-like episodic memory. Trends in Cognitive Sciences 29, 928–941 (2025).

[76] Spens, E. & Burgess, N. Hippocampo-neocortical interaction as compressive retrieval-augmented generation. Nature Communications 17, 7971 (2026).

[77] Schwartenbeck, P. et al. Generative replay underlies compositional inference in the hippocampal-prefrontal circuit. Cell 186, 4885–4897.e14 (2023).

[78] Sun, W., Advani, M., Spruston, N., Saxe, A. & Fitzgerald, J. E. Organizing memories for generalization in complementary learning systems. Nature Neuroscience 26, 1438–1448 (2023).

[79] Radford, A. et al. Language models are unsupervised multitask learners. OpenAI Blog (2019).

[80] Ho, J. et al. *Video diffusion models*. Advances in Neural Information Processing Systems, Vol. 35, 8633–8646 (2022).

[81] Michelmann, S., Kumar, M., Norman, K. A. & Toneva, M. Large language models can segment narrative events similarly to humans. Behavior Research Methods 57, 39 (2025).

[82] Kumar, M. et al. Bayesian surprise predicts human event segmentation in story listening. Cognitive Science 47, e13343 (2023).

[83] Schapiro, A. C., Turk-Browne, N. B., Botvinick, M. M. & Norman, K. A. Complementary learning systems within the hippocampus: A neural network modelling approach to reconciling episodic memory with statistical learning. Philosophical Transactions of the Royal Society B: Biological Sciences 372, 20160049 (2017).

[84] Zhu, J.-Y., Krähenbühl, P., Shechtman, E. & Efros, A. A. Generative visual manipulation on the natural image manifold. European Conference on Computer Vision (ECCV), 597–613 (2016).

[85] van der Maaten, L. & Hinton, G. Visualizing data using t-SNE. Journal of Machine Learning Research 9, 2579–2605 (2008).

[86] Ho, J., Jain, A. & Abbeel, P. Denoising diffusion probabilistic models. Advances in Neural Information Processing Systems, Vol. 33, 6840–6851 (2020).

[87] Ho, J. & Salimans, T. Classifier-free diffusion guidance. Preprint at 10.48550/arXiv.2207.12598 (2022).

[88] Simonyan, K. & Zisserman, A. Very deep convolutional networks for large-scale image recognition. International Conference on Learning Representations (2015).

[89] Kingma, D. P. & Ba, J. Adam: A method for stochastic optimization. International Conference on Learning Representations (2015).

[90] Ramsauer, H., et al. Hopfield networks is all you need. International Conference on Learning Representations (2021).

[91] Le, Y. & Yang, X. Tiny ImageNet visual recognition challenge. CS 231N, Stanford University (2015).

[92] van den Oord, A., Li, Y. & Vinyals, O. Representation learning with contrastive predictive coding. Preprint at 10.48550/arXiv.1807.03748 (2018).

[93] Hang, T., et al. Efficient diffusion training via Min-SNR weighting strategy. Proceedings of the IEEE/CVF International Conference on Computer Vision (ICCV), 7441–7451 (2023).

[94] Kay, K. N., Rokem, A., Winawer, J., Dougherty, R. F. & Wandell, B. A. GLMdenoise: A fast, automated technique for denoising task-based fMRI data. Frontiers in Neuroscience 7, 247 (2013).

[95] Virtanen, P. et al. SciPy 1.0: Fundamental algorithms for scientific computing in Python. Nature Methods 17, 261–272 (2020).

